# Human immune cells infiltrate the lesioned spinal cord and impair recovery after spinal cord injury in humanized mice

**DOI:** 10.1101/626721

**Authors:** Randall S. Carpenter, Roselyn R. Jiang, Faith H. Brennan, Jodie C.E. Hall, Manoj K. Gottipati, Stefan Niewiesk, Phillip G. Popovich

## Abstract

Immune compromised mice require ~4 months of engraftment with human umbilical cord blood CD34^+^ stem cells to develop a full and functional human immune system
The human neuroinflammatory response elicited after spinal cord injury in humanized mice is limited at 2 months post-engraftment but matures by 4 months
Intraspinal neuroinflammation consists of a florid human T cell and macrophage response, and human T cells co-localize with human macrophages
A human intraspinal neuroinflammatory response exacerbates lesion pathology and impairs functional recovery

**Abstract:** Humanized mice are a useful tool to help better understand how the human immune system responds to central nervous system (CNS) injury. However, the optimal parameters for using humanized mice in preclinical CNS injury models have not been established. Here, we show that it takes 3-4 months after engraftment of neonatal immune compromised mice with human umbilical cord stem cells to generate a robust human immune system. Indeed, sub-optimal human immune cell responses occurred when humanized mice received spinal contusion injuries at 2 months vs. 4 months post-engraftment. Human T cells directly contact human macrophages within the spinal cord lesion of these mice and the development of a mature human immune system was associated with worse lesion pathology and neurological recovery. Together, data in this report establish an optimal experimental framework for using humanized mice to help translate promising preclinical therapies for CNS injury.

## Introduction

Immunocompromised mice engrafted with human immune systems (i.e. “humanized” mice) are powerful pre-clinical models for studying human immune cell function. However, the translational value of these mice has not been fully realized, particularly in preclinical models of neuroinflammation and central nervous system (CNS) injury. Previously, we documented the feasibility of using humanized mice to study systemic and neuroinflammatory changes caused by traumatic spinal cord injury (SCI) (Carpenter et al., 2015). We did not explore developmental changes in the human immune systems of these mice. Specifically, the relative changes in the composition or function of the various human immune cells that could elicit neuroinflammation and systemic immunity after SCI. These are important variables to understand since there are conflicting data regarding the functional competency of human immune cells in humanized mouse models. Some studies indicate that human immune cells can develop in mice but they are not functional (Danner et al., 2011; Gille et al., 2012; Halkias et al., 2015; Strowig et al., 2010). Other published data indicate that both innate and adaptive human immune cells respond to inflammatory stimuli with robust effector functions (e.g., proliferation, cytokine production, antibody synthesis, migration toward chemotactic cues, etc.) (Cheng et al., 2017; Coughlan et al., 2012; Cravens et al., 2005; Ishikawa et al., 2005; Misharin et al., 2012; O’Boyle et al., 2012; Rodewohl et al., 2017; Shultz et al., 2010; Tanaka et al., 2012; Zayoud et al., 2013). Next-generation humanized mouse models with “improved” immune function are being generated to address these issues (Miller et al., 2013; Rathinam et al., 2011; Rongvaux et al., 2014; Saito et al., 2016; Takagi et al., 2012; Wetmore et al., 2012; Wunderlich et al., 2018). These conflicting reports could be explained by variability in the maturation state of human immune cells. Indeed, recent data indicate that human immune cell functions in humanized mice vary as a function of time post-engraftment (Audigé et al., 2017; Lang et al., 2013; Rodewohl et al., 2017). It takes time for undifferentiated human umbilical cord blood (UCB) stem cells to engraft bone marrow and secondary lymphoid tissues (e.g., spleen, lymph nodes) of immune compromised mice, differentiate into multi-lineage human progenitors, generate mature human immune cells, and then organize into immune niches. In immune competent mice, the development of the immune system occurs *in utero* (Holladay and Smialowicz, 2000).

In rodent SCI models, most experiments are performed in young adult animals at ~8-12 weeks of age. We predict that if humanized mice are injured at that age, as we have done previously (Carpenter et al., 2015), their human immune systems will be immature and functionally distinct from humanized mice in which longer periods of development were allowed after engraftment with human UCB stem cells. To test this hypothesis, we compared the composition and relative frequency of human PBLs as a function of time post-engraftment.

Here, we show that it takes ~3-4 months for human UCB stem cells to generate a robust human immune system in immune compromised mice. By 4 months, both human innate and adaptive immune cells respond to inflammatory stimuli, i.e., they form proliferative niches, produce antibodies and release cytokines indicating that they are functional. Importantly, when humanized mice receive an SCI at 4 months post-engraftment, as opposed to 2 months when the human immune system is still developing, notable differences in intraspinal inflammation were observed. Together, these novel data reveal an optimal experimental framework for using humanized mice to study human immune responses to CNS injury. These data also illustrate the adverse effects that post-injury activation of the human immune system can have on recovery of function.

## Results

### Human immune cell engraftment and composition is optimal by 4 months post-engraftment

To determine when peak development of a human immune system occurs in humanized mice, we quantified the percentage of human peripheral blood lymphocytes (hPBLs) up to 12 months after intrahepatic injection of human UCB CD34^+^ stem cells in NOD-SCID-IL2rγ^null^ (NSG) mouse pups (see Methods and also Carpenter et al., 2015; Huey and Niewiesk, 2018). At two months post-engraftment, human CD45^+^ (hCD45^+^) PBLs comprised 34% of total circulating leukocytes (**Fig. 1A**), increasing to 59% at 4 months post-engraftment. From 4-12 months post-engraftment, hPBLs were stable. In most hNSG mice, we achieved 30-90% human chimerism. Human chimerism was poor in only 5% (n=4/78) of hNSG mice, as defined by <10% hCD45^+^ PBLs at 4 months post-engraftment.

**Figure 1.**
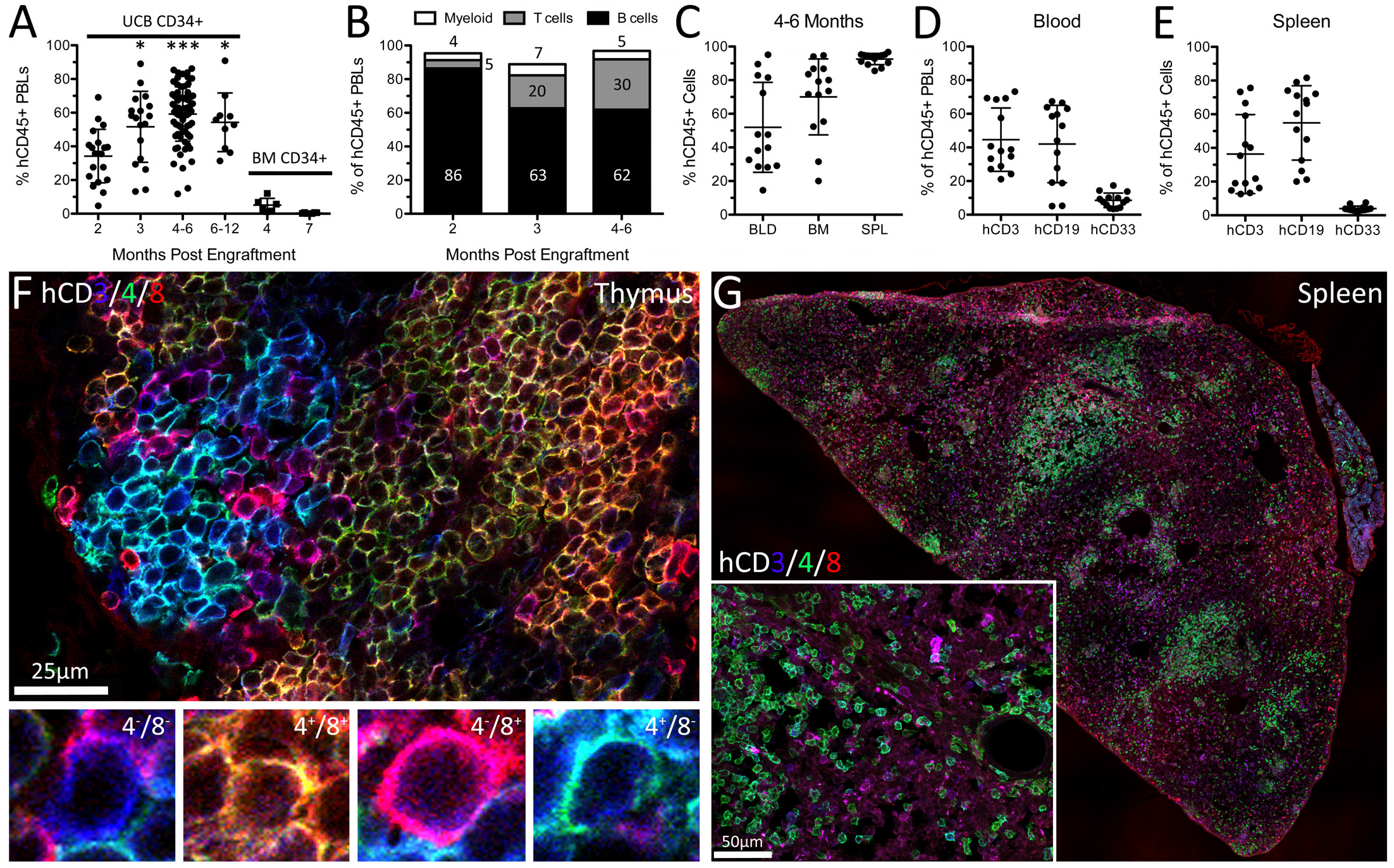
Long-lived engraftment of human immune systems in NSG mice occurs without toxicity. Human immune systems were derived from hCD34^+^ umbilical cord blood stem cells. A) Engraftment efficiency in mice is affected by time post-engraftment and source of stem cells. UCB = umbilical cord blood, BM = bone marrow. Data average ± SD; *p<0.05 ***p<0.001 compared to 2 months post-engraftment (UCB CD34^+^ only), oneway ANOVA with Tukey’s multiple comparison test. B) Proportion of human CD45^+^ peripheral blood leukocyte (PBL) subsets. C) Human CD45^+^ cells in the blood (BLD), bone marrow (BM), and spleen (SPL) of hNSG mice 4-6 months post-engraftment. D,E) Human immune subsets (hCD3, hCD19, and hCD33) in blood (D) and spleen (E) of hNSG mice 4-6 months post-engraftment. F,G) Human CD3^+^, CD4^+^, and CD8^+^ T cells identified in thymus (F) and spleen (G) of naïve hNSG mice 4 months post-engraftment. Human T cell subsets in thymus are highlighted (F).

In parallel with a plateau in the frequency of hPBLs, we noted a time-dependent change in the composition of circulating human immune cells (**Fig. 1B,D**). At 2 months post-engraftment, most human immune cells were CD19^+^ B cells (86%). At 4-6 months post-engraftment, the frequency of hCD19^+^ B cells decreased to 62%, concomitant with an increase in human CD3^+^ (hCD3^+^) T cells (30% of hPBLs). Human myeloid cells (hCD33^+^) comprised the smallest proportion of human cells from 2-6 months post-engraftment (4-7%).

By 4 months, when the frequency of circulating human PBLs reached a plateau, robust human immune cell engraftment was also observed in the bone marrow and spleen (**Fig. 1 C,E; Fig. 1-figure supplement 3A,B**). Human immune composition in the spleen matches the blood (**Fig. 1D,E**), with many human lymphocytes (T and B lymphocytes) and few myeloid cells. In the thymus of hNSG mice, mixed populations of hCD4^-^/hCD8^-^, hCD4^+^/hCD8^+^, hCD4^-^/hCD8^+^, and hCD4^+^/hCD8^-^ T cells could be found indicating ongoing thymopoiesis (**Fig. 1F**). NSG mice not injected with human stem cells had negligible thymus development. Spleens of humanized mice contained mostly human immune cells (**Fig. 1C**) with the white pulp densely populated by hCD3^+^/hCD4^+^ human T cells (**Fig. 1G**).

The source of human stem cells was also important. Compared to the robust chimerism that was achieved using UCB stem cells, adult human bone marrow CD34^+^ stem cells were inefficient as a donor source in the neonatal hNSG model. Only 1/6 mice developed >10% hCD45^+^ PBLs at 4 months post-engraftment with adult bone marrow stem cells and human chimerism was <1% in all mice by 7 months post-engraftment (**Fig. 1A**).

Collectively, these data illustrate that hNSG mice do not develop a mature human immune system until 3-4 months post-engraftment with human UCB CD34^+^ stem cells. This is an important biological variable that is not well-documented in the literature but if ignored, could dramatically affect experimental outcome (Audigé et al., 2017; Ishikawa et al., 2005; Lang et al., 2013; Rodewohl et al., 2017; Tanaka et al., 2012).

### Human immune cells respond to ex vivo and in vivo immune stimulation

To determine whether human immune cells in hNSG mice are functional at 4-6 months post-engraftment, human splenocytes were isolated, purified (see **Fig. 2-figure supplement 1A**) and then activated *ex vivo* using cell-specific stimuli. Human splenocytes were comprised mostly of hCD4^+^ T cells with lower numbers of hCD19^+^ B cells and hCD8^+^ T cells (**Fig. 2-figure supplement 1B**). In response to polyclonal stimulation with hCD3/28 and recombinant human IL2 (rhIL2), human T cells increased expression of hCD69 (**Fig. 2A,B**), a cell activation marker, accompanied by robust proliferation (**Fig. 2C,D; Fig. 2-figure supplement 1C**) and production of human IFNγ and IL-10 (**Fig. 2E,F**).

**Figure 2.**
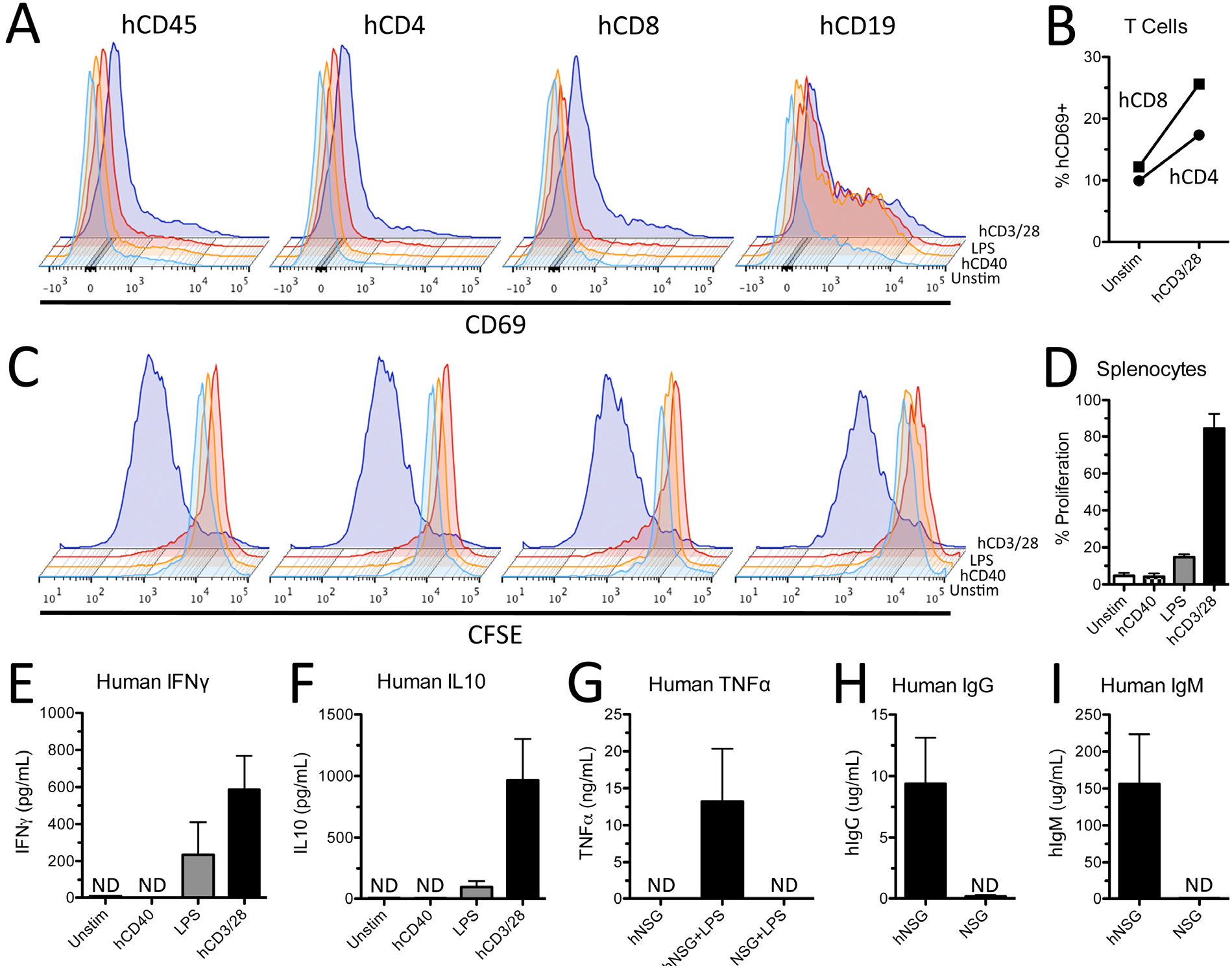
Human innate and adaptive immune cells from hNSG mice are functional and respond to cell-specific stimulation. A) Human splenocytes upregulate cell surface expression of activation marker CD69 48 hours after stimulation with human CD3/28 antibody and rhIL2. B) Proportion of hCD4^+^ and hCD8^+^ T cells expressing CD69 48 hours after stimulation by hCD3/28 and rhIL2. C) Decrease in CFSE staining demonstrating robust proliferation of human splenocytes stimulated with hCD3/28 and rhIL2. D) Proportion of proliferating splenocytes 96 hours after cell specific stimulation. E,F) Quantification of human interferon gamma (IFNγ) and IL10 in culture supernatants after 96 hours of cell specific stimulation. G) Human TNFα quantification in blood serum 1 hour after *in vivo* injection with 3 mg/kg lipopolysaccharide (LPS). Human IgG (H) and IgM (I) from blood serum in hNSG mice. Note the absence of human cytokines and antibodies in blood serum of non-engrafted NSG mice treated with LPS, demonstrating species specificity of ELISAs. ND=not detected. Data average ± SEM; n=2 biological replicates in B,D; n=4 biological replicates in E,F; n=3 mice per group in G,H; n=3 NSG and n=6 hNSG mice in I,J.

When stimulated with a hCD40 activating antibody (clone 5C3) and rhIL4, purified human B cells increased hCD69 expression (**Fig. 2A,B**) but did not proliferate or produce cytokines (**Fig. 2C-F**). However, stimulation of human T cells with hCD3/28 and rhIL2 induced robust hCD19^+^ B cell proliferation (**Fig. 2C**), suggesting human T and B cell interactions are necessary for mediating human B cell function.

Lipopolysaccharide (LPS), a canonical activator of toll-like receptor 4 (TLR4) found mostly on myeloid cells, also increased proliferation and production of human cytokines by human splenocytes (**Fig. 2C-F**). Similarly, LPS injected *in vivo* (3mg/kg, i.p.) elicited production of human TNFα (hTNFα) by 1-hour post-injection (**Fig. 2G**). Human TNFα was not detected in serum from naive hNSG mice or non-humanized NSG mice injected with LPS. We also detected human IgG and IgM antibodies in blood serum of hNSG mice but not non-humanized NSG mice (**Fig. 2H,I**). Together, these data prove that human immune cells from hNSG mice are functional; they respond *ex vivo* and *in vivo* to physiologically-relevant stimuli, producing a range of immune effector molecules (e.g., cytokines, antibodies).

### Time post-engraftment determines human peripheral immune cell responses to contusive SCI

Data in **Figs. 1–2**, together with published data (Audigé et al., 2017; Lang et al., 2013; Rodewohl et al., 2017), indicate that the composition, relative density and functional maturity of human immune cells increases as a function of time post-engraftment in humanized mice. Thus, one would predict that the effects of SCI on immune system activation and the subsequent effects of neuroinflammation and lesion histopathology will change as a function of time post-engraftment. To test this hypothesis, a single human UCB donor was used to generate two cohorts of hNSG mice born 2 months apart. Both groups received a SCI at the same time – one at 2 months post-engraftment and the other at 4 months post-engraftment. Both cohorts survived to 35 days post-injury (dpi). Consistent with data from **Fig. 1**, uninjured hNSG mice had more human PBLs at 4 months post-engraftment than at 2 months (**Fig. 3A; Fig. 3-figure supplement 1A**) – all circulating human leukocyte subsets including T lymphocytes (hCD3), B lymphocytes (hCD19) and myeloid cells (hCD33) increased in number and proportion as a function of time post-engraftment (**Fig. 3B; Fig. 3-figure supplement 1B**).

**Figure 3.**
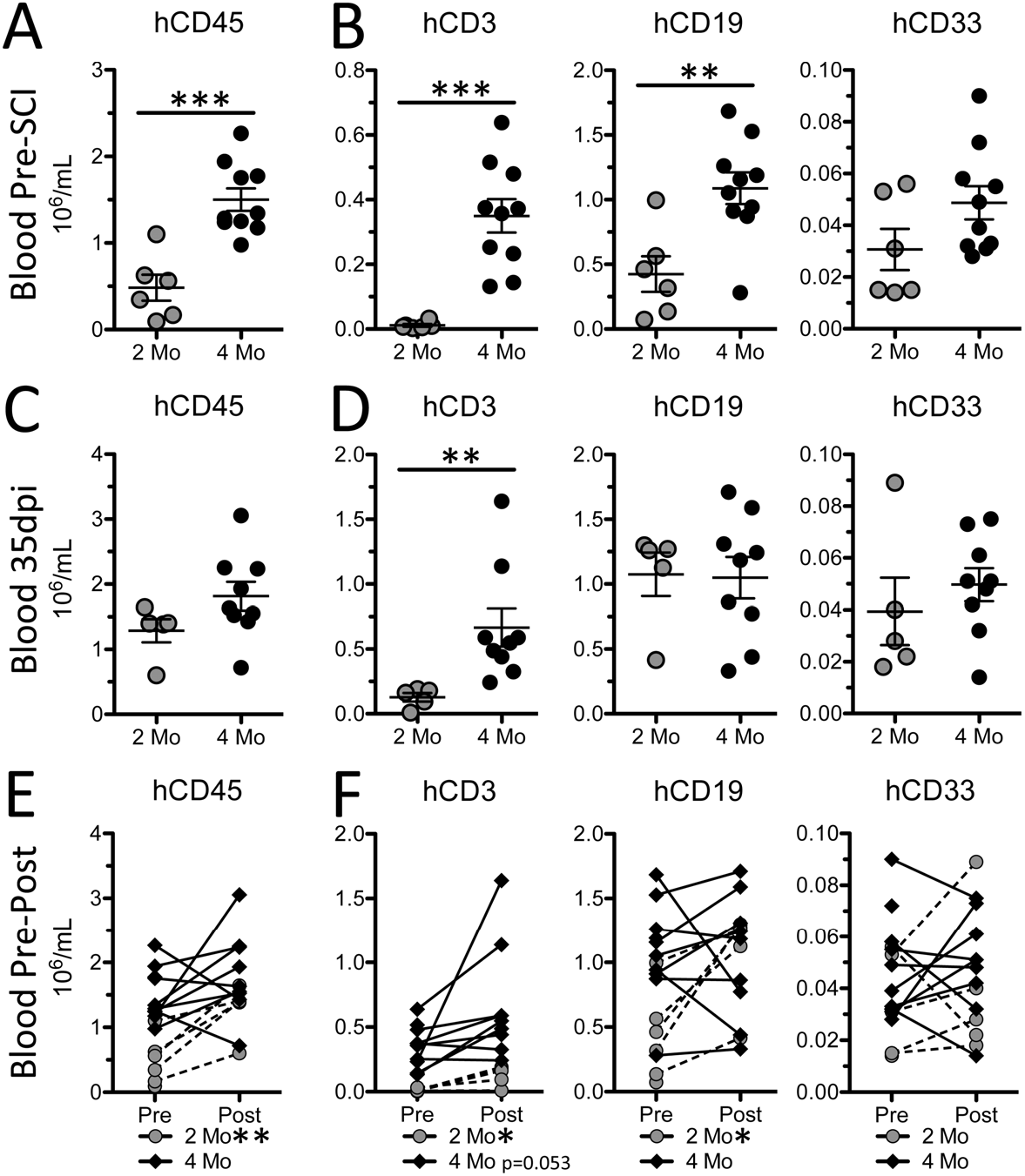
Human peripheral blood leukocyte responses to SCI differ as a function of time post-engraftment. Total numbers of human PBLs (A,C), and human PBL subsets (B,D), 7 days prior to and 35 days after SCI. Change in numbers of human PBLs (E), and human PBL subsets, from pre-to post-SCI. *p<0.05 **p<0.01 ***p<0.001 student’s unpaired (A-D) and paired (E,F) t-test. Data average ± SEM.

Five weeks after SCI we noted marked differences in circulating leukocyte responses between the two cohorts. In the 2-month cohort, the total numbers of circulating human leukocytes increased after SCI (**Fig. 3E,F; Fig. 3-figure supplement 1E,F**). However, this was unlikely an injury-dependent effect and is better explained by continued maturation of the human immune system. Indeed, by 35 dpi cells from mice in the 2-month cohort were now engrafted >3 months with the relative proportion of hCD45^+^ PBLs in most mice increased to levels identical to engraftment at 3 months (**compare Fig. 1A,B**) and similar to pre-injury values found in the 4-month post-engraftment cohort (**compare Fig. 3A,C; Fig. 3-figure supplement 1A,C**). Consistent with this interpretation, post-injury changes in hCD45^+^ PBLs were modest in the 4-month post-engraftment cohort (**Fig. 4E**) and were marked by a selective increase in the proportion of T cells (**Fig. 3F; Fig. 3-figure supplement 1F**).

**Figure 4.**
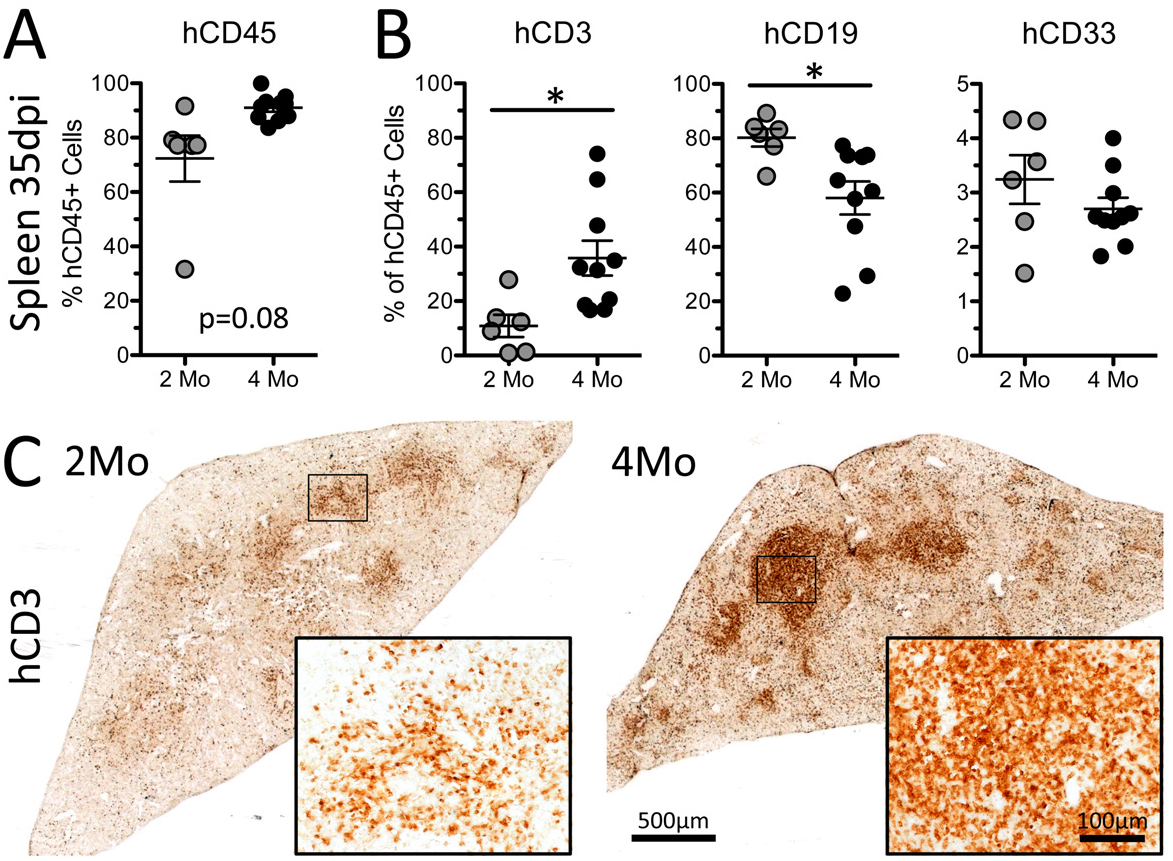
Human splenocyte proportion and composition 35 days after SCI in hNSG mice differ as a function of time post-engraftment. Human splenocyte chimerism (A) and composition (B) at 35 dpi between hNSG mice injured at 2- and 4-months post-engraftment. C) Human CD3^+^ T cells in spleens of hNSG cohorts. Images are representative of the mean hCD3 values for both groups. *p<0.05 student’s t-test. Data average ± SEM.

Given the importance of the spleen in regulating neuro-immune interactions after SCI (Blomster et al., 2013; Noble et al., 2018), we also quantified human splenocyte subsets in both cohorts 35 dpi (**Fig. 4A,B**). The relative proportions of human T, B, and myeloid cells in spleen reflected their proportions in the blood 35 dpi, which is consistent with the role of the spleen as a reservoir for circulating human leukocytes. Spleens of both groups displayed preferential clustering of hCD3^+^ T cells in the splenic white pulp. Again, the relative density of T cell clusters was increased in the 4-month post-engraftment cohort (**Fig. 4C**).

### Time post-engraftment determines the magnitude and composition of human T cell infiltration into the injured spinal cord

Previously, we showed that SCI in humanized mice elicits neuroinflammation in the injured spinal cord, comprised mostly of mouse microglia, human macrophages and human lymphocytes (Carpenter et al., 2015). To determine whether time post-engraftment affects the magnitude of human immune cell recruitment and the ability of these cells to influence spinal cord anatomy, we compared recruitment of human leukocytes in the 2- and 4-month cohorts. Infiltration by hCD45^+^ leukocytes was increased in SCI mice from the 4-month cohort (**Fig. 5A,B**). Additionally, hCD3^+^ T cells were increased ~10-fold in mice injured at 4 months post-engraftment compared to 2 months post-engraftment (**Fig. 5C,D**).

**Figure 5.**
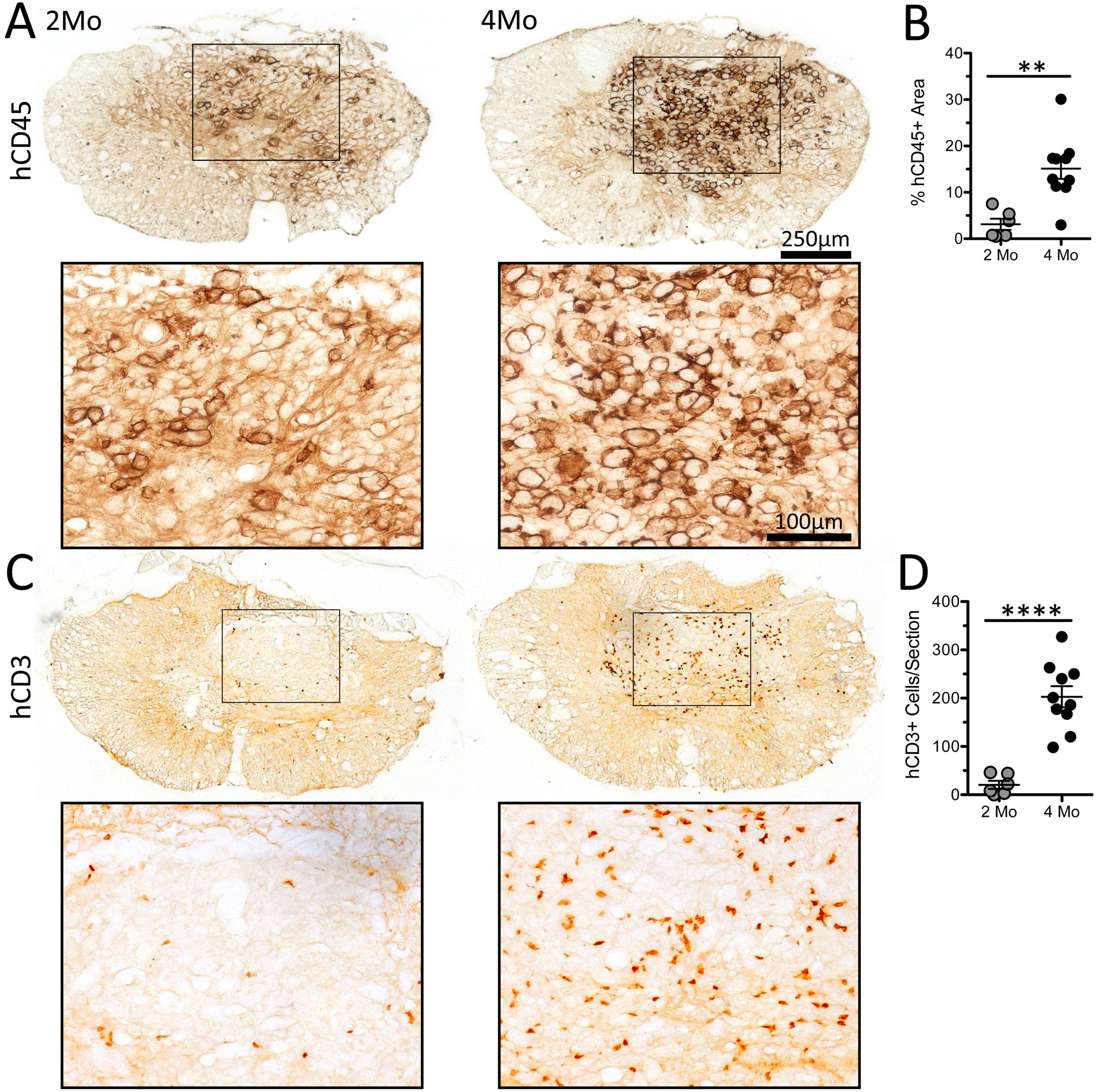
Time post-engraftment determines the magnitude of human immune cell infiltration into the injured spinal cord of hNSG mice. A) Human CD45^+^ leukocytes within the lesion epicenter, primarily confined to the central lesion. B) Proportional area analysis of hCD45^+^ immunolabeling within the lesion. C) Human CD3^+^ T cells within the lesion epicenter, with quantification (D). Images representative of the mean for each data set. **p<0.01 ****p<0.0001 student’s t-test. Data average ± SEM.

Multi-color immunofluorescent confocal microscopy identified hCD3^+^/hCD4^+^ helper and hCD3^+^/hCD8^+^ cytotoxic T cell subsets within spinal cord lesions (**Fig. 6A**). We never observed colocalization of hCD4^+^ and hCD8^+^ labeling on any single hCD3 T cell. In line with blood and spleen data in **Figs. 4–5**, more human T cell subsets were consistently found in the injured spinal cord of hNSG mice from the 4-month cohort (**Fig. 6B**). The composition of helper to cytotoxic T cells in spinal cord lesions of hNSG mice injured at 4 months post-engraftment (70%:30%) was significantly different than at 2 months (50%:50%) (**Fig. 7C**).

**Figure 6.**
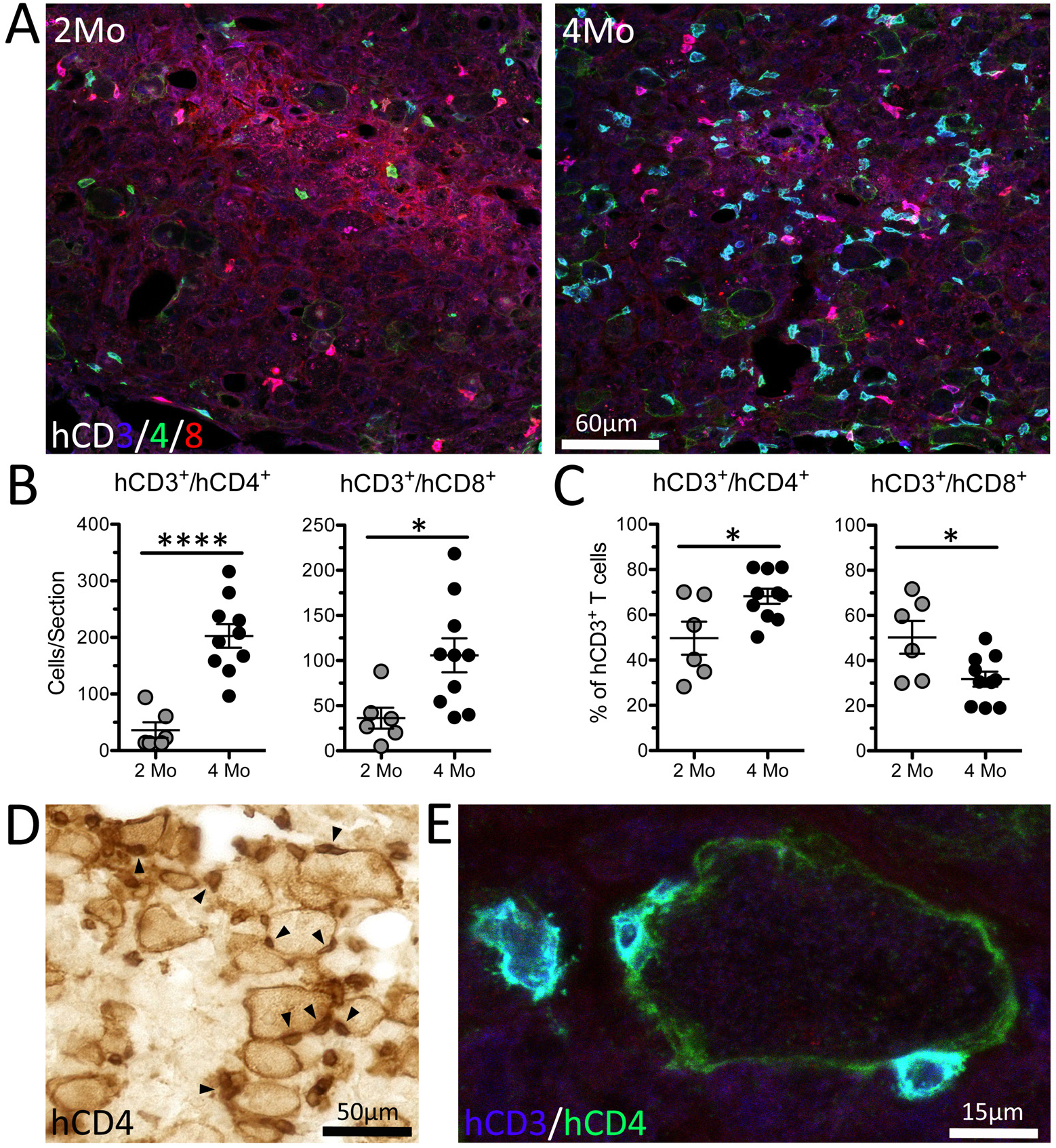
Human helper and cytotoxic T cell subsets infiltrate the injured spinal cord, with T helper subsets directly contacting human macrophages. A) Immunofluorescent labeling of hCD3^+^/hCD4^+^ helper and hCD3^+^/hCD8^+^ cytotoxic T cells at the lesion epicenter 35 dpi in 2- and 4-month post-engraftment hNSG mice. MIPAR automated quantification of total numbers (B) and relative proportion (C) of human T cell subsets. D) Immunolabeling of hCD4 identifies both small and large cells with morphology of T cells and macrophages, respectively. Small hCD4^+^ T cells were often found directly adjacent to large, phagocytic human macrophages, indicating T cell-macrophage interaction within the injured spinal cord (arrowheads). E) Confocal imaging of hCD3^+^/CD4^+^ T cells directly contacting a hCD3^-^/hCD4^+^ macrophage. *p<0.05 ****p<0.0001 student’s t-test. Data average ± SEM.

**Figure 7.**
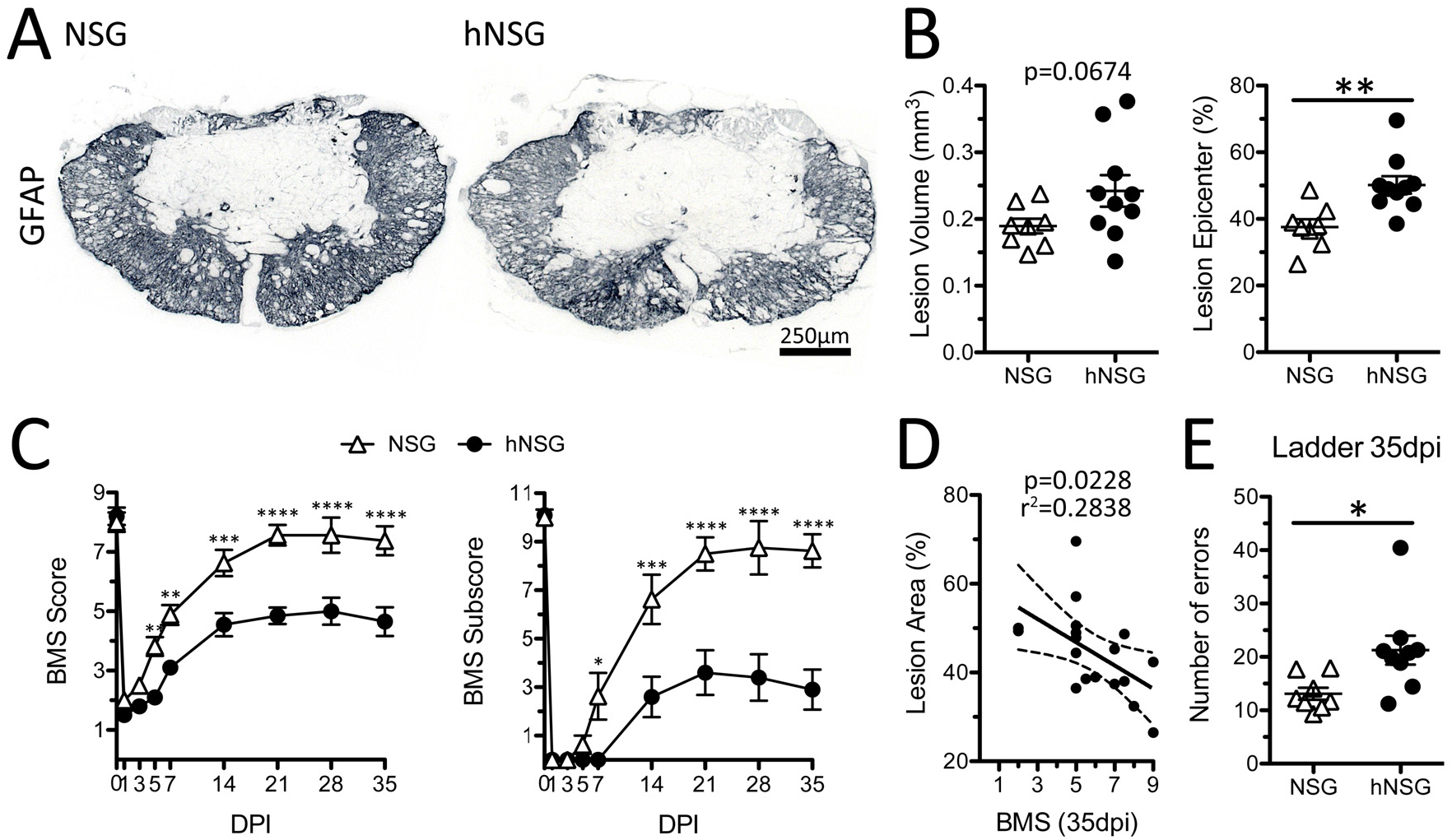
Humanized NSG mice (4-months post-engraftment) have increased lesion pathology and impaired functional recovery after SCI compared to age-matched non-engrafted NSG mice. A) Astrocyte (GFAP) labeling of injury epicenter. B) Lesion volume and area (proportion) at epicenter as defined as GFAP-negative region. C) BMS score and subscore from 1-35 dpi. D) Linear regression analysis of lesion size and BMS scores at 35 dpi. E) Number of errors (foot falls) on a horizontal ladder test 35 dpi. *p<0.05 **p<0.01 ***p<0.001 ****p<0.0001 student’s t-test (B,F) and repeated measures two-way ANOVA with Bonferroni multiple comparisons test (C). Data average ± SEM.

We’ve previously shown that large, phagocytic human macrophages populate the injured spinal cord in humanized mice (Carpenter et al., 2015). The spinal cord lesions of both hNSG cohorts contained a subset of large (presumably activated phagocytic) human macrophages expressing hCD4 on their cell membrane (**Fig. 6D,E**). Although CD4 selectively labels helper T lymphocytes in mice, both CD4 and CD8 are expressed by human and rat macrophages (Boddaert et al., 2018; Crocker et al., 1987; Gibbings and Befus, 2009; Gibbings et al., 2007; Hiraki et al., 2009; Ikeda et al., 2014; Popovich et al., 2003; Sroga et al., 2003; Szabo et al., 1990). In the injury epicenter, human helper T cells (hCD3^+^/hCD4^+^) were often found in direct contact with hCD4^+^ macrophages (arrowheads, **Fig. 6D,E**). These T cell-macrophage interactions were found in both hNSG cohorts, and while not directly quantified, more T cell-macrophage interactions were found in the 4-month cohort. Together, data in **Figs. 5&6** indicate that recruitment of human leukocytes to the lesion epicenter increases in mature humanized mice, creating neuroinflammatory lesions that are indistinguishable from what occurs after SCI in conventional mice, rats and humans (Beck et al., 2010; Fleming et al., 2006; Jones et al., 2005, 2004, 2002; Kigerl et al., 2006; Sroga et al., 2003; Wu et al., 2017).

### A human immune system in NSG mice worsens lesion pathology and hindlimb functional recovery after SCI

Post-injury recruitment of T lymphocytes is a regular component of the neuroinflammatory reaction in mouse, rat and human (Beck et al., 2010; Fleming et al., 2006; Jones et al., 2005, 2004, 2002; Kigerl et al., 2006; Sroga et al., 2003; Wu et al., 2017). In pre-clinical mouse and rat SCI models, both injurious and protective effects have been attributed to infiltrating T cells after SCI (E. Hauben et al., 2000; Ehud Hauben et al., 2000; Jones, 2014; Jones et al., 2005, 2004, 2002; Liu et al., 2019; Raposo et al., 2014; Satzer et al., 2015; Schwartz, 2001; Schwartz and Kipnis, 2001; Schwartz and Moalem, 2001; Sun et al., 2017). For obvious reasons, establishing a causal role for infiltrating T cells in either injury or repair has not been possible in people with SCI.

Previously, we found that lesion pathology was exacerbated and functional recovery was impaired in hNSG mice as compared to non-engrafted immunocompromised NSG control mice (Carpenter et al., 2015). However, our previous study occurred before we understood the importance of post-engraftment timing on maturation of the human immune system hNSG mice. Here, we compared lesion pathology and functional recovery in non-engrafted NSG mice and age-matched hNSG mice with a mature (4-month post-engraftment) human immune system after spinal contusion injury. hNSG mice develop larger lesions (**Fig. 7A,B**) with significantly worse recovery of hindlimb function as defined by open field locomotor analysis (**Fig. 7C**) and foot-falls on the horizontal ladder test (**Fig. 7E**). Worse hindlimb function at 35 dpi correlated with an increase in lesion size (**Fig. 7D**). These data indicate that post-injury activation of human immune cells, like mouse immune cells, can exacerbate lesion pathology and impair neurological recovery after SCI (Carpenter et al., 2015).

## Discussion

Previously, we characterized the anatomical, functional and hematological consequences of a contusive spinal cord injury (SCI) in humanized NSG (hNSG) mice (Carpenter et al., 2015). Although the data in that report were the first to illustrate the feasibility of using humanized mice to test hypotheses related to neuro-immune interactions after SCI, the report was limited in scope and did not evaluate the implications of developmental timing on human immune cell composition and function.

Data in this report show the relative efficiency of human chimerism and corresponding changes in human immune cell composition change as a function of time post-engraftment. Human chimerism peaks between 3-4 months post-engraftment and remains stable for at least 12 months. Of the human immune cells that we characterized, T lymphocytes are likely key to the development of a mature human immune system in hNSG mice and may ultimately play a key role in influencing post-injury neuroinflammation, pathology and recovery of function.

In humanized mice, human T cells help promote the maturation and functional development of human B cells (Lang et al., 2013). In athymic nude rats, which lack T cells, B cells are similarly impaired but can be rescued by reconstituting adult rats with T cells (Milićević et al., 2005). It is not clear how T cells influence the development and maturation of other immune cells but interactions between T cells and progenitor cells in the bone marrow seem to be essential (Russell et al., 2015; Schürch et al., 2014).

Our data indicate that it takes ~4 months post-engraftment before T cells reach a stable plateau in the blood of hNSG mice. This delay in human T cell development could explain why some studies, which typically used hNSG mice at earlier post-engraftment times, have concluded that human immune cells in hNSG mice are not functional (Danner et al., 2011; Gille et al., 2012; Halkias et al., 2015; Strowig et al., 2010). By 4 months post-engraftment, we found that innate and adaptive human immune cells were functionally competent; i.e., they responded *ex vivo* and *in vivo* to various physiologically-relevant stimuli, producing a range of immune effector molecules (e.g., cytokines, antibodies). These data are consistent with other data showing that inflammatory stimuli elicit robust effector functions in human immune cells from humanized mice (e.g., cytokine production, antibody synthesis, migration toward chemotactic cues, etc.) (Cheng et al., 2017; Coughlan et al., 2012; Cravens et al., 2005; Ishikawa et al., 2005; Misharin et al., 2012; O’Boyle et al., 2012; Rodewohl et al., 2017; Shultz et al., 2010; Tanaka et al., 2012; Zayoud et al., 2013). Although we did not measure human immune cell function at earlier post-engraftment times, we can infer from our comparative SCI studies that human immune system development and function are sub-optimal in hNSG mice at 2 months post-engraftment. Specifically, the magnitude of the neuroinflammatory response after SCI, as defined by numbers of intraspinal hCD45^+^ leukocytes and human T cells (hCD3, hCD4, hCD8), was markedly reduced in hNSG mice injured at 2 months post-engraftment compared to 4 months post-engraftment (Figs. 5&6). Notably, the magnitude of the human T cell infiltrate and relative proportions of hCD4^+^ and hCD8^+^ subsets in the injured spinal cord of hNSG mice in the 4 month cohort was consistent with what is observed in conventional C57BL/6 and C57BL/10 mice after SCI (Kigerl et al., 2006; Sroga et al., 2003).

More human macrophages were also found in spinal cord lesions in the 4-month cohort (as compared with lesions from hNSG mice in the 2-month cohort). A more robust myeloid cell response to SCI was observed even though the percentage of human myeloid cells in blood or spleen of hNSG mice was not different between mice in the 2- and 4-month post-engraftment cohorts. Again, this is evidence that a more functionally mature human immune system exists in hNSG mice at 4 months post-engraftment.

We also found that many large, phagocytic human macrophages expressed membrane hCD4. Rat CD4^+^ macrophages are found at sites of myocarditis (Baba et al., 2006), in tumors (Ikeda et al., 2014) and in the injured rat spinal cord (Popovich et al., 2003; Sroga et al., 2003). Some CD4^+^ macrophages also have been identified in the mouse gut and thymus, with unique ontogeny and function (Esashi et al., 2003; Zangerle-Murray et al., 2018). This co-receptor, which is routinely used to phenotype helper T-cells, is also expressed by human and rat monocytes (Crocker et al., 1987; Mestas and Hughes, 2004). Ligating this co-receptor on monocytes/macrophages triggers intracellular signaling cascades that augment macrophage maturation and activation (Szabo et al., 1990, Zhen et al., 2014). Although the functional significance of CD4^+^ macrophages in CNS trauma has not been determined, macrophage expression of CD8, another co-receptor typically found on cytotoxic T cells, has been linked to pathological effector functions and pathology in the rat models of multiple sclerosis (Hiraki et al., 2009), stroke (Boddaert et al., 2018) and spinal cord injury (Popovich et al., 2003). Therefore, humanized mice could prove to be a useful tool for studying the functional importance of activating CD4 (and possibly CD8) co-receptors on CNS macrophages.

Using confocal microscopy, we confirmed several examples of cell-cell contact between human T cells and human macrophages in the injured spinal cord. Membrane hCD3 expression was abundant at the interface between human macrophages and human T cells (see **Fig. 6-figure supplement 2**). Redistribution of CD3, a major component of the T cell receptor (TCR) complex, is essential in the formation of immune synapses and T cell activation by macrophages and other antigen presenting cells (Fooksman et al., 2010). We did not have a means to evaluate intraspinal T cell activation in hNSG mice; however, mouse and rat macrophages and T cells have been shown to have injurious and neuroprotective effects after SCI (Ankeny et al., 2009; Brennan and Popovich, 2018; Gensel et al., 2009; Goldstein et al., 2016; Ehud Hauben et al., 2000; Jones et al., 2004; Kigerl et al., 2009; Popovich et al., 2002; Raposo et al., 2014; Schwartz and Kipnis, 2001; Shechter and Schwartz, 2013; Walsh et al., 2015, 2014).

In the present report, data suggest that the intraspinal human immune response contributes to pathology and subsequent functional impairment. Indeed, after SCI in 4-month post-engraftment hNSG mice with mature functional human immune systems, a prominent intraspinal human T cell infiltrate was associated with exacerbated lesion pathology and impaired spontaneous recovery. Still, we previously provided evidence that axons migrate along lesion matrix proteins in close proximity to human immune cells in hNSG mice, indicating possible reparative functions of human immune cells. Together, these data demonstrate that in hNSG mice, the human immune system actively responds to SCI hNSG mice and that these cells exert biological effects on the lesion epicenter. To prove a causal role for human T cell-mediated pathology or repair, human immune cell manipulation studies are needed.

### Considerations for using humanized mice in CNS injury models

Long-term health and viability of humanized mice may be a concern for those interested in brain or spinal cord aging studies or if chronic survival times are required. We were consistently able to keep hNSG mice alive for 9-12 months of age without major health concerns. Sporadic premature death of hNSG mice does occur but cause of death is difficult to determine. Anemia may be one factor contributing to premature mortality (Knibbe-Hollinger et al., 2015; Rongvaux et al., 2014; Yoshihara et al., 2019). Progressive destruction of mouse RBCs can occur with high levels of human engraftment (Knibbe-Hollinger et al., 2015; Rongvaux et al., 2014; Yoshihara et al., 2019) and new transgenic humanized mice with improved engraftment and maturation of human innate immune cells typically die 3-4 months after engraftment because of rapid onset anemia (Rongvaux et al., 2014). SCI in hNSG did not increase the frequency of death, although we did not allow survival beyond 35 days post-SCI. hNSG mice developed infrequent health problems (e.g., ruffled fur, cataracts in one or both eyes and spontaneous hindlimb inflammation). Importantly, SCI did not increase the number or severity of these health issues. We did note that hNSG mice appear healthier when they are engrafted with human UCB stem cells close to 72 hours after birth. This anecdotal observation requires empirical testing.

A criticism of humanized mice is that they do not produce robust human immune responses *in vivo* (Danner et al., 2011; Gille et al., 2012; Halkias et al., 2015; Strowig et al., 2010). As mentioned above, the belief that humanized mice are immunologically impaired has been perceived as a weakness of the model but in reality, when assays are performed at later post-engraftment intervals, human immune cells respond as expected. In our hands, a stable human immune system requires at least 3-4 months to develop in hNSG mice. The development of a functionally competent human immune system may also be affected by other factors that vary between labs including strain of the recipient immune deficient mice, age of engraftment (newborn vs. young adult mice), route of human stem cells injection (intraperitoneal vs. intravenous vs. intrahepatic), human stem cell source (UCB vs. fetal liver vs. adult bone marrow), stem cell phenotype (cell surface expression of CD34, CD38 and CD133), stem cell purity, stem cell dose, supplements given during development (e.g., rhIL7), engraftment of additional human tissues (fetal liver/thymus), etc. Each of these variables should be considered when interpreting published data; however, none are disqualifying factors that would preclude the use of humanized mice in experimental context of CNS trauma.

Although there are various types of humanized mouse models, including several that include “next generation” strategies for manipulating the composition and function of the human immune system in mice (see Shultz et al., 2012, 2011, 2007 for review), we used the hNSG model because it is the current standard across a range of scientific disciplines (Audigé et al., 2017; Dykstra et al., 2016; Lang et al., 2013; Rodewohl et al., 2017; Tanaka et al., 2012).

Moreover, hNSG mice are available through commercial vendors, although current costs may be prohibitive for larger studies (~$1,000 USD/mouse). Fortunately, it is possible to reduce costs and generate hNSG mice “in house”. In our experience, it is possible to generate a litter of 8-10 hNSG mice for ~$1,500 USD – that cost includes the purchase and housing of breeding pairs, a single vial of human UCB stem cells (each vial contains ~5×10^5^ cells) and housing per-diem costs for ~6 months. The final per mouse cost is ~$150-200 USD. Generating your own hNSG mice also make it possible for sex to be incorporated as a biological variable in the design of experiments (Clayton, 2016); commercial vendors only provide ready to use cohorts of female mice.

### Conclusion

Data in the current report indicate that optimal human immune responses require ~4 months of development in hNSG mice. Once activated by SCI, human immune cells infiltrate the injured spinal cord where, through unknown mechanisms, they contribute to pathology and impair recovery of function. Using data and guidelines established in this report, future research can incorporate humanized mice into experimental plans to better understand how SCI, or other types of neurological injury or disease, affect human immune system function *in vivo*. In this context, humanized mice provide a unique step in the translational research spectrum and should facilitate the study of human immune cell-mediated mechanisms and testing of human-specific interventions *in vivo* before moving therapies into human clinical trials.

## Methods

### Humanized mice

The Institutional Animal Care and Use Committee of The Ohio State University approved all animal protocols. Adult male and female NOD.Cg-Prkdc^scid^IL2rg^tm1Wjl^/SzJ (NSG mice) breeding pairs were purchased from The Jackson Laboratory (strain #005557) and bred in-house. Animals were fed commercial food pellets and chlorinated reverse osmosis water ad libitum. Mice were housed ≤5/cage in ventilated microisolator cages containing with corn cob bedding, and the housing facility in a 12-hour light-dark cycle at a constant temperature (20 ± 2°C) and humidity (50 ± 20%). NSG strain mice were kept in sterile housing conditions during breeding and throughout the humanization process. Generation of NSG mice with human immune systems (hNSG) was performed as previously described (Carpenter et al., 2015; Huey and Niewiesk, 2018; Pearson et al., 2008). Briefly, 24-72 hour postnatal NSG pups received 100cGy whole body irradiation (RS 2000, Rad Source, Suwanee, GA), followed immediately by engrafting 1-5×10^4^ human umbilical cord CD34^+^ stem cells (Lonza Incorporated, Walkersville, MD; or Stemcell Technologies, Vancouver, BC) via intrahepatic injection. After injection pups were allowed to recover for several minutes on a heat source set to 37°C before returning pups to their dams for normal maturation and weaning at 21-24 days of age. hNSG mice are healthy and can be used over extended periods of time for experimentation. On occasion, hNSG mice can have a rough coat and decreased body weight compared to non-humanized NSG mice. Cataract formation in one or both eyes was also observed in a few animals, and 1/78 hNSG mice developed spontaneous hindlimb inflammation around the knee. Median survival of our initial cohort of hNSG mice was 293 days. While death was preceded by rapid weight loss over ~1 week, there were no phenotypic changes associated with classical graft-versus-host disease (GvHD), such as loss of fur and advanced infiltration of cells into organs and tissues. Only 4% (3/78) of hNSG mice displayed evidence of T-cell expansion (excessive thymus size or proportion of T cells in peripheral blood), further supporting a lack of graft-versus-host disease (GvHD) in this model. Both male and female hNSG mice generated robust human immune systems. The efficiency and composition of engrafted human immune cells differed only slightly between male and female NSG mice in blood at 4 months post-engraftment. Total engraftment by human immune cells was 7.5% higher in females (63.0 ± 14.2% vs. 55.5 ± 17.2%), with more B cells (67.2 ± 16.3% vs. 56.2 ± 17.6%) and fewer myeloid cells (3.91 ± 2.8% vs. 6.73 ± 3.8%). Overall, these data support the use of both male and female humanized mice in experimental studies.

### SCI & animal care

hNSG mice (2-4 months old) were used for SCI experiments. hNSG mice used for comparison of time post-engraftment were generated 2 months apart using same UCB donor but underwent SCI and all other outcomes at the same time. Mice were anesthetized with ketamine (90-120mg/kg, i.p.) and xylazine (7-10mg/kg, i.p.) then injected with gentamicin sulfate (5mg/kg, s.q.). Aseptic conditions were maintained during all surgical procedures. Mice were placed on a warming pad during surgery to maintain body temperature. Hair was shaved at the region of the thoracic spinal cord and skin treated with a sequence of betadine-70% ethanol-betadine. A small midline incision was made to expose the mid-thoracic vertebra, then a partial laminectomy was performed. A 60 KDyne (mild) T9 contusion injury was performed using an Infinite Horizon Impactor (Precision Systems and Instrumentation, Lexington, KY). Immediately after impact, the muscle overlaying the injury site was sutured, followed by closure of the wound with sutures or staples. After surgery, mice were placed in cages on heating pads and monitored until they recovered consciousness. Animals were supplemented with sterile saline (1-2mL, s.q.) and softened food to eat *ad libitum* during recovery. Bladders were expressed twice daily, and urine underwent weekly pH testing to detect bladder infections. Gentocin antibiotic was subcutaneously administered once a day at 5 mg/kg for 5 days post-injury (dpi).

### Hindlimb locomotor function assessment

The open-field Basso Mouse Scale (BMS) was used to assess hindlimb paralysis and functional recovery (Basso et al., 2006). Pre-SCI testing occurred after acclimating mice to the open field on multiple days with dim lighting to minimize anxiety. Post-SCI testing occurred on 1, 3, 5, 7, 14, 21, 28, and 35 dpi. Mice explored the open field and were scored over a period of 4 minutes. Two individuals that were blinded to the experimental condition averaged the left and right hindlimb score for each mouse to obtain a single score per mouse at each time point.

The horizontal ladder test was performed before SCI and at 35 dpi. The horizontal ladder contained 33 rungs spaced ~1 cm apart and elevated ~1 cm off of a clear mirrored surface for viewing of paws from the side and below using a digital video recorder to capture each trial. Mice were acclimated to the ladder test over 3 separate days with a minimum of 3 runs per session before acquiring baseline values. Mice were handled by a single experimenter (RSC). During testing days, each mouse was allowed a minimum of 3 passes along the horizontal ladder to reduce trial-trial variability. Mice were allowed to rest for ~15-20 minutes between each pass/trial. Digital videos were assessed by a single experimenter (RRJ) blinded to group designations, with playback at 0.35x speed using VLC media player (v.3.0; VideoLAN Organization) on a computer screen of a minimum 13-inch diameter. Each trial (3/mouse) was viewed and analyzed 3 times by the experimenter and averaged to acquire a single value for each trial. This was done to reduce observer error and variability, and a subset of trials were assessed by a second reviewer (RSC) to ensure accuracy.

### Tissue collection

For survival blood sampling, blood was collected from the submandibular vein using a lancet and EDTA-coated capillary tube (Sarstedt Inc., Thermo Fisher Scientific, Waltham, MA). For endpoint tissue collection, mice were first deeply anesthetized with ketamine and xylazine. Blood was collected via cardiac puncture, placed in EDTA-coated collection tubes, and inverted several times. Red blood cells were lysed with ammonium chloride and resuspended in 0.1M phosphate buffered saline (PBS) with 2% FBS (flow buffer) for flow cytometry. Spleens were isolated, weighed, and placed in Hank’s Balanced Salt Solution (HBSS). Spleens were minced with sterile dissection scissors and smashed through a 40 μm sterile filter using the plunger of a 3 mL syringe and 10 mL of HBSS. Mouse femurs and tibiae were removed, cleaned, and placed in a small volume of HBSS. Bone marrow cells were isolated by either flushing bones with 10 mL HBSS or by crushing in a mortar and pestle with HBSS. Cell counts for bone marrow and spleen were obtained by standard hemocytometer counting techniques, while total blood was analyzed with a Hemavet 950fs multi-species hematology system capable of analyzing whole blood cells with 5-part differential, platelets, and red blood cells. To obtain serum, blood was collected as above, inserted into gel separation tubes coated with clotting activator, and centrifuged at 10^5^ x g for 5 minutes. For experiments using tissue sections, mice were perfused with 0.1M PBS followed by 4% paraformaldehyde. Fixed tissues (i.e., the spleen, thymus and spinal cords) were cryopreserved for 48 hours in 30% sucrose, cryosectioned at 10 μm onto glass slides, and stored at −20°C until staining.

### Immunolabeling and imaging

Antibody sources, working concentrations, protocols, and validations were previously reported (Carpenter et al., 2015); a summary of antibodies for immunolabeling are included in Table 1. Slides were thawed on a slide warmer and then rinsed in 0.1M PBS. Endogenous peroxidases were quenched with 6% H2O2 diluted in methanol for 15 minutes at room temperature. Sections were incubated with 4% BSA, 3% normal goat serum, and 0.1% Triton X-100 in 0.1 M PBS for 1 hour followed by primary antibody overnight at 4°C. Secondary antibody incubation was performed at room temperature for 1 hour. Slides were then incubated with Vectastain ABC (Vector Laboratories, Burlingame, CA) for 1 hour at room temperature and developed with 3,3’-diaminobenzidine (DAB/ImmPACT DAB) or Vector SG (Vector Laboratories, Burlingame, CA) of for 5-15 minutes (until optimal differentiation of signal). A separate protocol was used to reduce non-specific labeling of mouse antibodies on mouse tissue to identify human immune cell types. After thawing, tissue sections were placed in ice-cold methanol for 15-20 minutes, followed by washes in PBS. When required (i.e. hCD3, hCD8, hCD4), antigen retrieval was accomplished by incubating tissues in heated alkaline pH Tris-based solution (Vector Laboratories) followed by a blocking step using a mixture of 4% BSA, 3% normal goat serum and 0.1% Triton X-100 in 0.1 M PBS for 1 hour. Sections were incubated with primary antibodies overnight at 4°C, followed by washes in PBS and subsequent incubation at room temperature for 1 hour with Alexa Fluor-conjugated secondary antibodies. Immunolabeled tissues were imaged using an Axioplan 2 imaging microscope equipped with an AxioCam digital camera and AxioVision v.4.8.2 software (Carl Zeiss Microscopy GmbH, Jena, Germany), or on a Leica TCS SCP confocal microscope (Leica Microsystems Inc., Buffalo Grove, IL).

**Table 1.** Antibodies

| *Antigen* | *Host, dilution* | *RRID* | *Vendor, catalog number* |
| --- | --- | --- | --- |
| **Primary antibodies** |  |  |  |
| **GFAP** | Rabbit, 1:5000 | AB_10013382 | Agilent, Z0334 |
| **hCD45** | Mouse, 1:250 | AB_400305 | BD Biosciences, 347460 |
| **hCD3** | Rat, 1:250 | AB_321245 | Bio-Rad, MCA1477 |
| **hCD4** | Rabbit, 1:500 | AB_2686917 | Abcam, ab183685 |
| **hCD8** | Mouse, 1:250 | AB_395995 | BD Biosciences, 555631 |
| **Secondary amplification/detection** |  |  |  |
| **Biotin-rabbit IgG** | Goat, 1:1000 | AB_2313606 | Vector Labs, BA-1000 |
| **Biotin-rat IgG** | Rabbit, 1:1000 | AB_10015300 | Vector Labs, BA-4001 |
| **Alexa488-rabbit IgG** | Goat, 1:500 | AB_2576217 | Invitrogen, A-11034 |
| **Alexa546-mouse IgG** | Goat, 1:500 | AB_144695 | Invitrogen, A-11030 |
| **Alexa633-rat IgG** | Goat, 1:500 | AB_141553 | Invitrogen, A-21094 |
| **Flow cytometry antibodies (per test)** |  |  |  |
| **mFc Block, CD16/32** | Rat, 1 μL | AB_394657 | BD Biosciences, 553142 |
| **hFc Block** | Unknown, 1 μL | AB_2728082 | BD Biosciences, 564219 |
| **mCD45-APC** | Rat, 2.5 μL | AB_398672 | BD Biosciences, 559864 |
| **hCD45-PerCP** | Mouse, 10 μL | AB_400307 | BD Biosciences, 347464 |
| **hCD3-APC-H7** | Mouse, 4 μL | AB_1645731 | BD Biosciences, 641397 |
| **hCD3-Pe-Cy7** | Mouse, 5 μL | AB_396896 | BD Biosciences, 557851 |
| **hCD4-APC** | Mouse, 4 μL | AB_314082 | BioLegend, 300514 |
| **hCD8-Pe-Cy7** | Mouse, 4 μL | AB_314130 | BioLegend, 301012 |
| **hCD19-BV510** | Mouse, 5 μL | AB_2737912 | BD Biosciences, 562947 |
| **hCD33-FITC** | Mouse, 10 μL | AB_400047 | BD Biosciences, 340533 |
| **hCD69-BV421** | Mouse, 3 μL | AB_10933255 | BioLegend, 310929 |

### Lesion pathology

Glial Fibrillary Acidic Protein (GFAP) labeled sections were imaged using a 5x objective. Images were loaded into ImageJ, and a random dot overlay (each dot representing 0.01 mm^2^) was placed on the image using the grid feature. Total number of dots per section, and number of dots within the GFAP-negative region, was determined and the proportion of the section area and total lesion volume calculated. Human CD3^+^ T cells within the lesions of hNSG mice were manually quantified on an Axioplan 2 microscope with a 20x objective. Two sections at the lesion epicenter (100 μm apart) were quantified, and values were averaged for each animal. Images of T cell subsets (hCD3^+^/hCD4^+^ and hCD3^+^/hCD8^+^) were acquired with a Leica TCS SCP confocal microscope with 20x magnification. Tiling with automatic stitching generated a high-resolution image covering the whole section, and imaging parameters were kept consistent to minimize variability. Z stacks (2 μm steps) were compressed to create one maximum intensity projection (MIP) image. Because T cells predominantly home to the gray matter (Kigerl et al., 2006), and the spared white matter contained high background staining, only spinal lesion and gray matter was included in the analysis. First, a mask of the lesioned gray matter was created using the freehand selection tool in ImageJ. Masks of the lesioned gray matter were imported into MIPAR software (Sosa et al., 2014). T cells were counted using custom cell counting algorithms (available upon request). Briefly, for quantification of human T helper cells (hCD3^+^/hCD4^+^), each channel was selected to generate two layers and a basic threshold for staining intensity was applied to each layer. Cells >50 pixels in size were removed from each layer and touching cells were split. Cells meeting stain and size criteria with spatially overlapping channels counted as one helper T cell. Batch processing applied this algorithm to all images, and the cell count data was exported to Microsoft Excel. This method was repeated for counting of human cytotoxic (hCD3^+^/hCD8^+^) T cells.

### Flow cytometry

1×10^6^ bone marrow cells and splenocytes, or approximately 50μL RBC-lysed blood, were allocated for flow cytometry analysis. Human and mouse Fc receptors were blocked for 15 minutes, followed by labeling with antibodies against specific antigens for 1 hour. A summary of antibodies and amounts added to samples for flow cytometry are included in Table 1. Dead cells were labeled with eFluor780 (eBioscience, Thermo Fisher Scientific) approximately 30 minutes into antibody incubation. Labeled cells were fixed and permeabilized with BD Fix/Perm solution (BD Biosciences, Franklin Lakes, NJ) for 20 min. All incubations were performed at 4°C and followed by a wash with excess flow buffer and centrifugation for 5 minutes at 4°C. An LSR II flow cytometer (BD Biosciences) analyzed immunolabeled cell samples. Forward scatter and side scatter parameters gated viable cell populations for phenotypic analysis using antibody panels. Offline data analyses were completed with FlowJo v.10 software (Tree Star, Inc., Ashland, OR).

### In vivo lipopolysaccharide challenge

hNSG and non-engrafted NSG mice (4-6 months old) were injected 3 mg/kg (i.p.) with lipopolysaccharide (LPS from *E. coli;* O55:B5, Sigma-Aldrich) or 0.9% saline solution. Mice were anesthetized, blood collected for serum, and tissues placed in 4% PFA overnight at 4°C. Blood was collected into a BD Microtainer serum separator tube with clotting activator, and after 30 minutes tubes were centrifuged at 10,000xg for 5 minutes. Serum was collected, aliquoted into 1.5mL Eppendorf tubes, and stored at −80°C.

### Ex vivo purification and stimulation of human splenocytes

hNSG mice (4-6 months old) were anesthetized and spleens dissected as previously described. Splenocytes were diluted in IMDM and counted on a manual hemocytometer with a 1:1 ratio of 0.4% Trypan Blue to quantify live cells. Contaminating mouse splenocytes were depleted from cell preparations by incubating with MACS anti-mouse CD45 magnetic microbeads (Miltenyi Biotec, Auburn, CA) and then washing through magnetic LD Columns as per manufacturer’s protocol. Enrichment of human cells was verified via flow cytometry of the depleted and cultured fractions: depleted fraction contained >90% mouse CD45^+^ splenocytes, while the cultured fraction contained >90% human CD45^+^ splenocytes (see **Fig. 2-figure supplement 1**).

Prior to culture and stimulation, human splenocytes were stained with CFSE using CellTrace CFSE Cell Proliferation Kit (Invitrogen) as per manufacturer recommendations. Human splenocytes were then cultured at 1×10^6^ cell per mL in ImmunoCult-XF T cell Expansion Medium (Stemcell Technologies, Vancouver, BC). For T cell stimulation, ImmunoCult Human CD3/CD28 T cell Activator and 20ng rhIL2 (Stemcell Technologies) was added to culture medium. For B cell stimulation, anti-human CD40 antibody (clone 5C3) and 20 ng rhIL4 (Stemcell Technologies) was added to culture medium. For LPS stimulation, 1 μg/mL was added to culture medium. Splenocyte cultures were maintained at 37°C in a 95% air and 5% CO2 incubator for 48-96 hours.

### Quantification of cytokines and antibodies from blood serum and culture supernatants

Human TNFα was quantified in blood serum (1:10 dilution) using a DuoSet ELISA kit (R&D Systems, Minneapolis, MN). Human IFNγ and IL10 were quantified in culture supernatants (1:2 dilution) using DuoSet ELISA kits (R&D Systems). DuoSet ELISA Ancillary Reagent Kit 2 was used in conjunction with all DuoSet ELISA kits. Human IgG and IgM was quantified in blood serum (1:1000 dilution) and culture supernatants (DILUTION) using the Human IgG/IgM total ELISA Ready-SET-Go! Kits (Invitrogen/eBioscience). All ELISAs were performed as per manufacturer’s protocols. Plates were read at 450nm wavelength on a SpectraMAX190 and analyzed using the SoftMax Pro software (Molecular Devices, San Jose, CA).

### Statistical analysis

GraphPad Prism software (version 5.0, La Jolla, CA) was used for data analysis. Statistical significance was assigned to a p< 0.05. Statistical tests are reported in figure legends and include unpaired t-test (with or without Welch’s correction for unequal variance), one-way ANOVA (with Tukey’s post-hoc test), repeated-measures two-way ANOVA (with Bonferroni post-hoc test), and Kaplan-Meier survival curve estimation. Data are represented as mean ± standard error of the mean except where noted. One sample from both 2- and 4-month post-engraftment groups was not run on the Hemavet due to inadequate blood volume at 35 dpi and was excluded from analysis in Fig. 4C&D. Adobe Photoshop CS5 v.12 (Adobe Systems Inc., San Jose, CA) was used to generate figures.

## Acknowledgements

This study was supported in part by Craig H. Neilsen Foundation Grant 340884 (PGP) and Fellowship 457267 (FHB), NIH Fellowship F31NS100303 (RSC), and the Ray W. Poppleton Endowment (PGP). Funding agencies were not involved in the study design, collection and interpretation of data, or preparation of the manuscript. The authors would like to thank Zhen Guan, Feng Qin Yin, and Vishal Kalbavi for their technical assistance. Flow cytometry data in this report were collected using instruments and services at the Analytical Cytometry Core, The Ohio State University, supported by NCI P30CA016058.

## Figure Supplements

**Figure 1-figure supplement 1.**
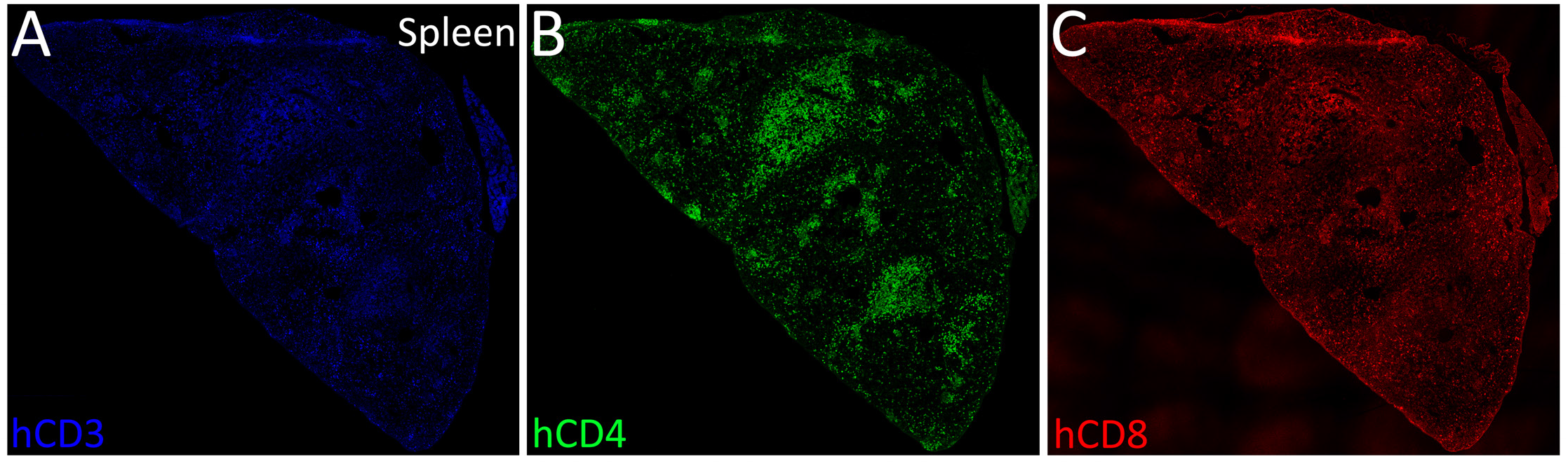
Isolated fluorescent channels from low-magnification confocal image used in Figure 1E demonstrating human CD3 (A), human CD4 (B), and human CD8 (C) immunolabeling in the spleen of a naïve hNSG mouse 4-months post-engraftment.

**Figure 1-figure supplement 2.**
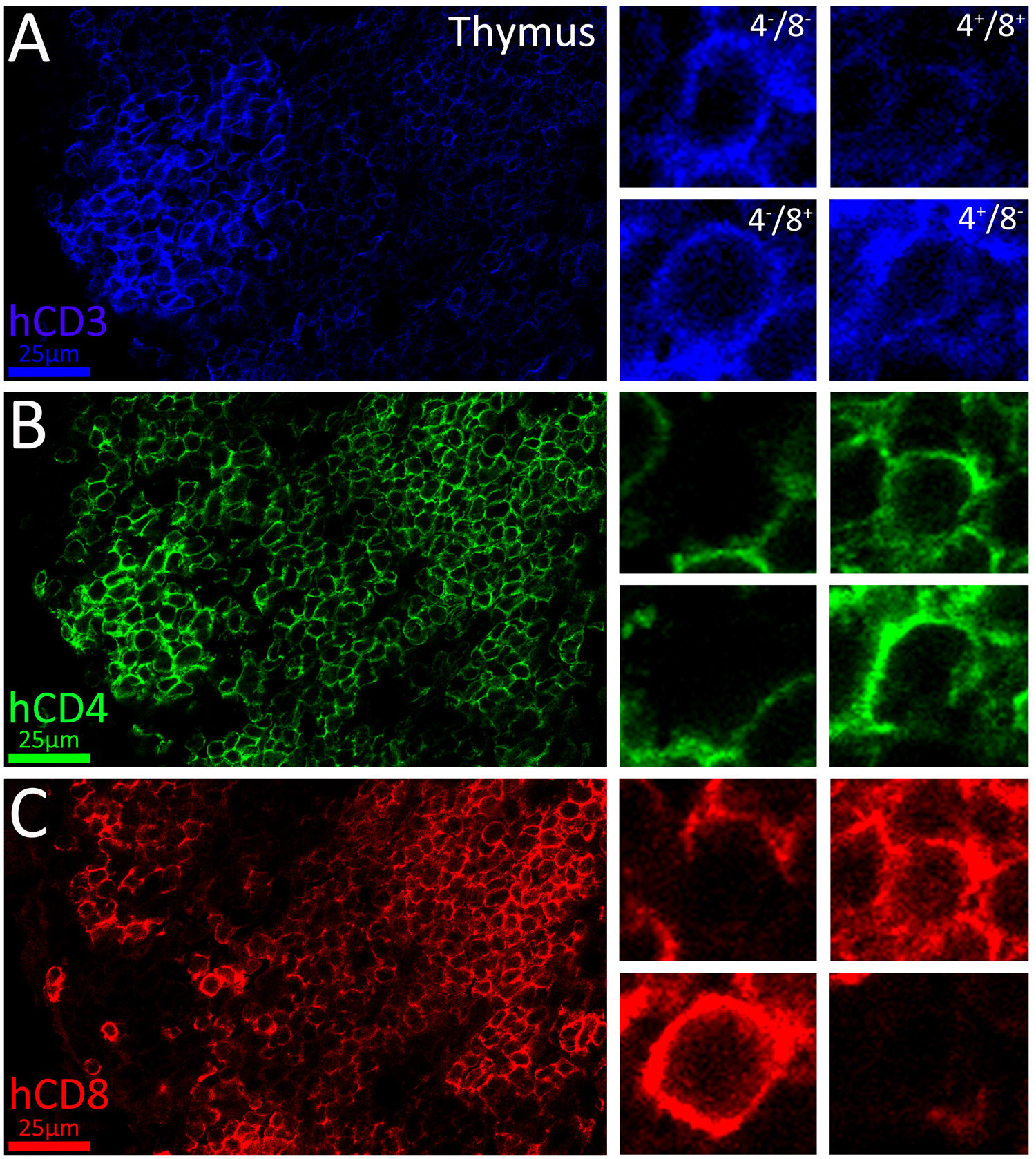
Isolated fluorescent channels from confocal images used in Figure 1F demonstrating human CD3 (A), human CD4 (B), and human CD8 (C) immunolabeling in the thymus of a naïve hNSG mouse 4-months post-engraftment. Human T cell subsets in thymus are highlighted with high-resolution confocal microscopy.

**Figure 1-figure supplement 3.**
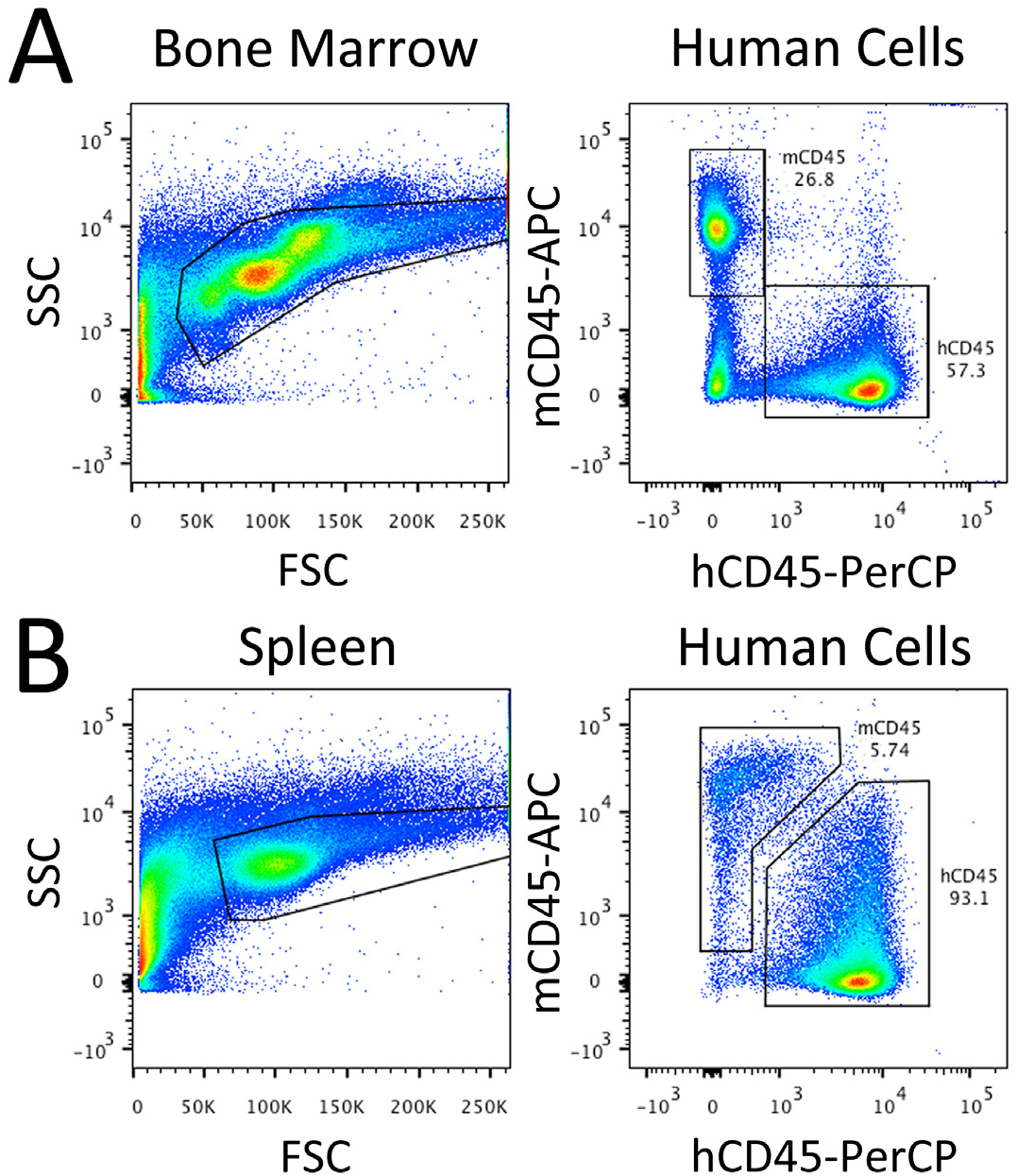
Example flow cytometry plots identifying human CD45^+^ immune cells in bone marrow (C) and spleen (D) of a naïve hNSG mouse 4 months post-engraftment. Left plots demonstrate gating of cells by size (FSC = forward scatter) and complexity (SSC = side scatter. Right plots demonstrate gating of mouse and human CD45^+^ immune cells using fluorescently labeled antibodies.

**Figure 2-figure supplement 1.**
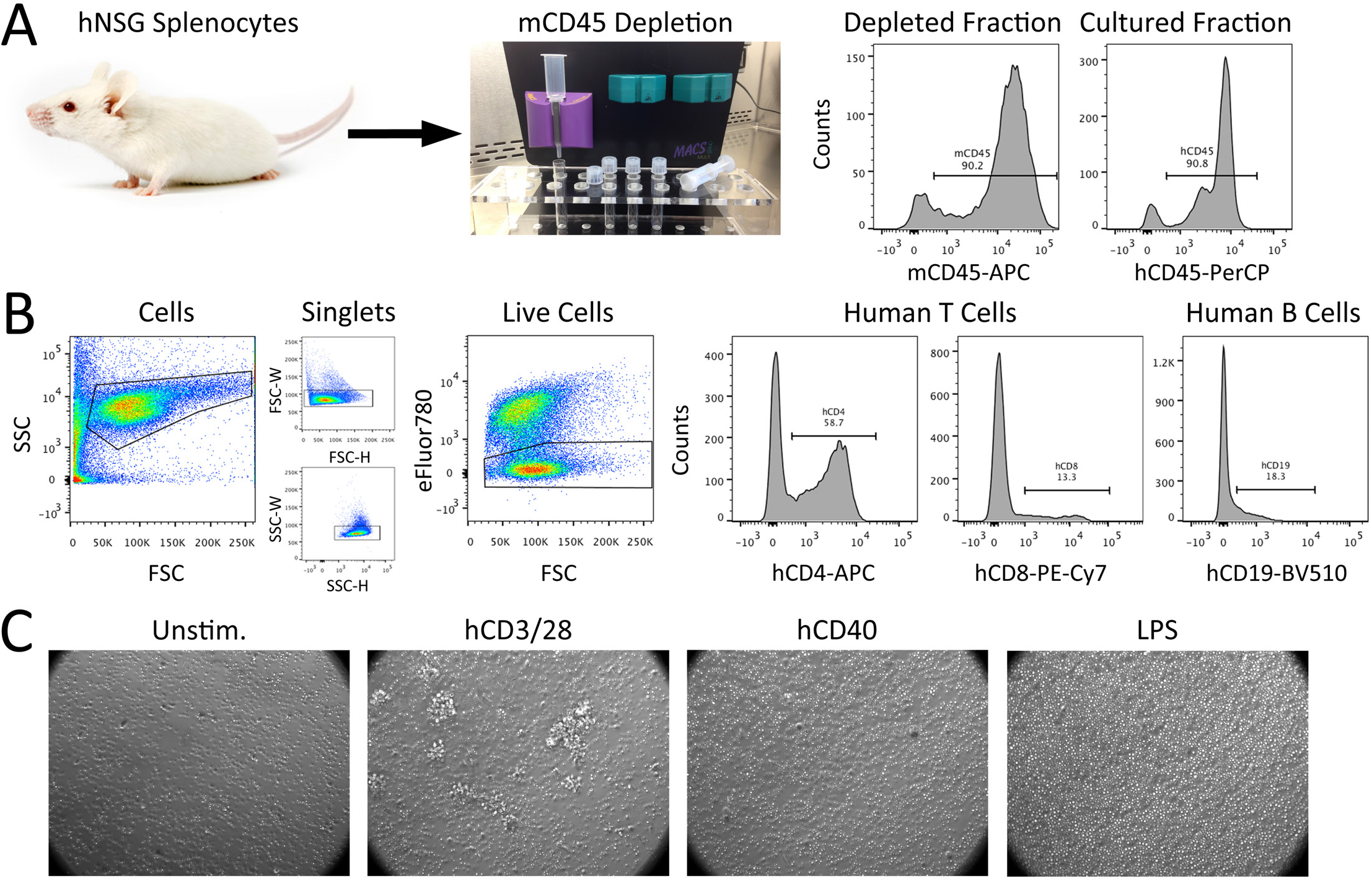
Isolation and flow cytometric analysis of human splenocytes from hNSG mice. A) Mouse splenocytes were depleted from hNSG splenocyte preparations using Miltenyi MACS magnetic bead depletion of mouse CD45^+^ cells. Flow cytometry confirmed mouse splenocyte depletion, with >90% of depleted cells expressing mouse CD45, and >90% of cultured cells expressing human CD45. Human splenocytes were then left unstimulated (unstim) or stimulated with either lipopolysaccharide (LPS), human CD40 monoclonal antibody with rhIL4, or human CD3/28 antibody with rhIL2. B) Example flow cytometry analysis of single cells after 48-96 hours in culture, including hCD4^+^ and hCD8^+^ T cells, and hCD19^+^ B cells. C) Low-magnification images of culture conditions after 48 hours. Note the appearance of proliferating cell clusters in the hCD3/28 culture well.

**Figure 3-figure supplement 1.**
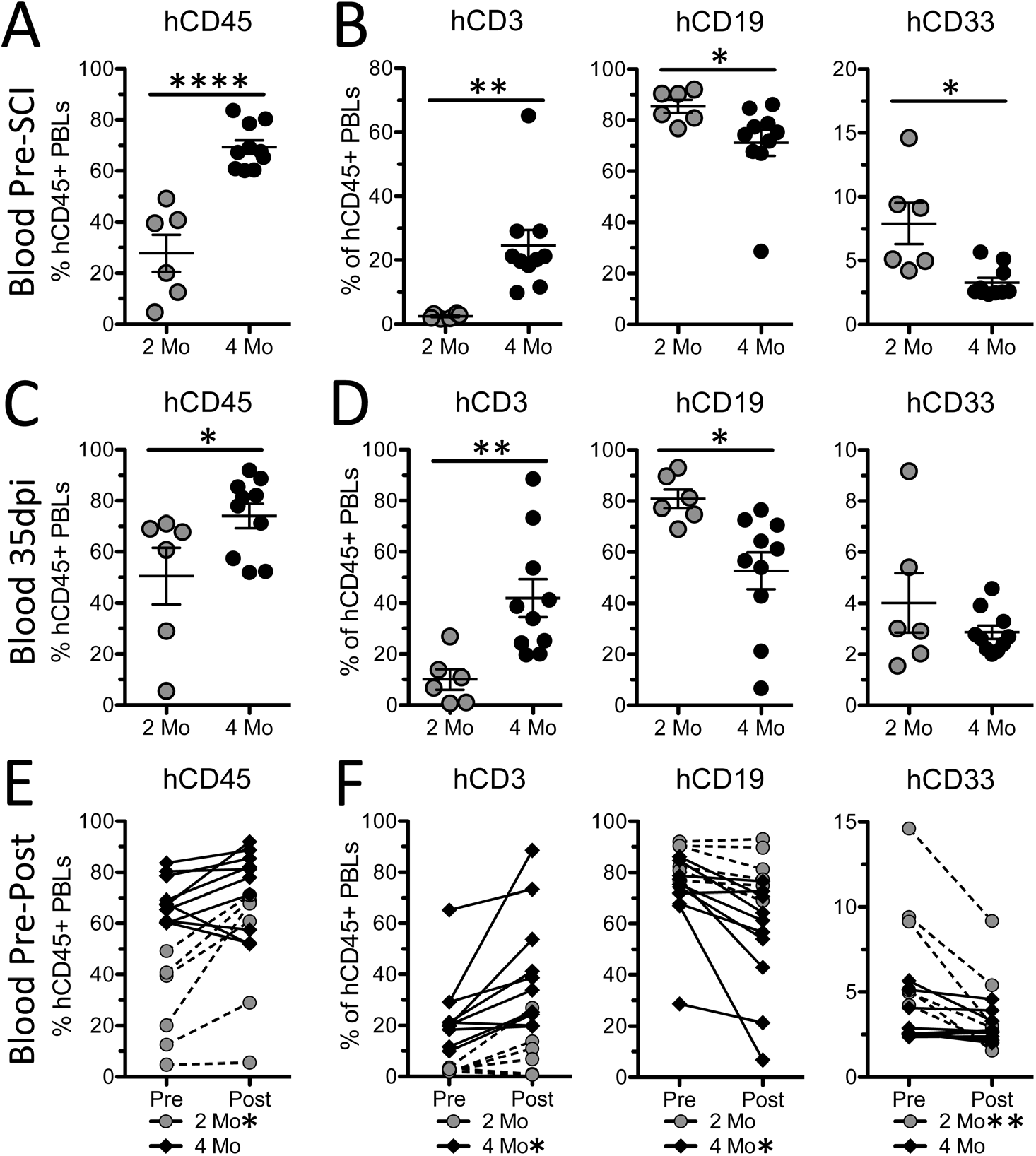
Human peripheral blood leukocyte responses to SCI differ as a function of time post-engraftment. Proportion of human PBLs (A,C), and human PBL subsets (B,D), 7 days prior to and 35 days after SCI. Change in proportion of human PBLs (E), and human PBL subsets, from pre-to post-SCI. *p<0.05 **p<0.01 ****p<0.0001 student’s unpaired (A-D) and paired (E,F) t-test. Data average ± SEM.

**Figure 6-figure supplement 1.**
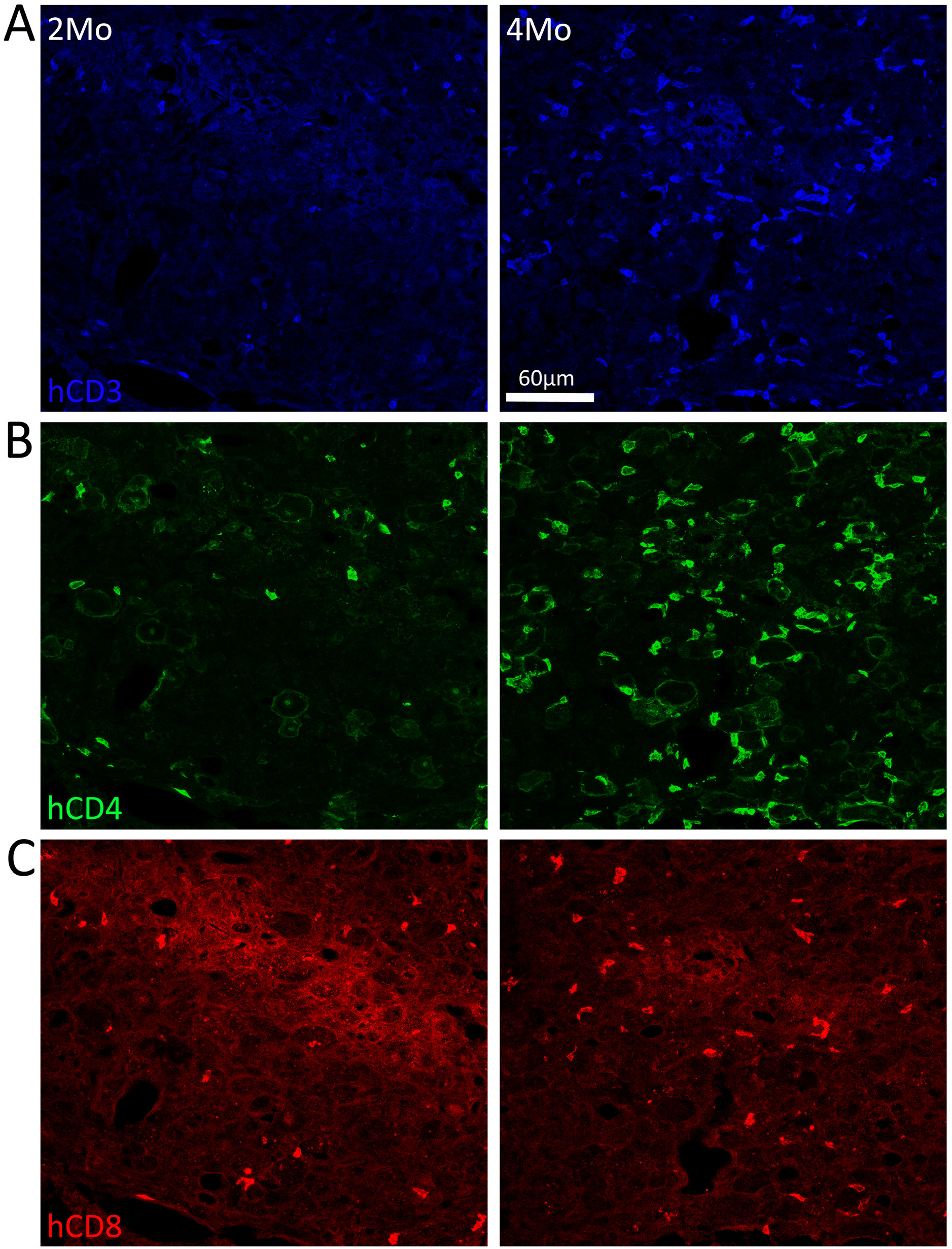
Isolated fluorescent channels from confocal images used in Figure 6A demonstrating human CD3 (A), human CD4 (B), and human CD8 (C) immunolabeling in lesion epicenters 35 dpi in hNSG mice injured at either 2- or 4-months post-engraftment.

**Figure 6-figure supplement 2.**
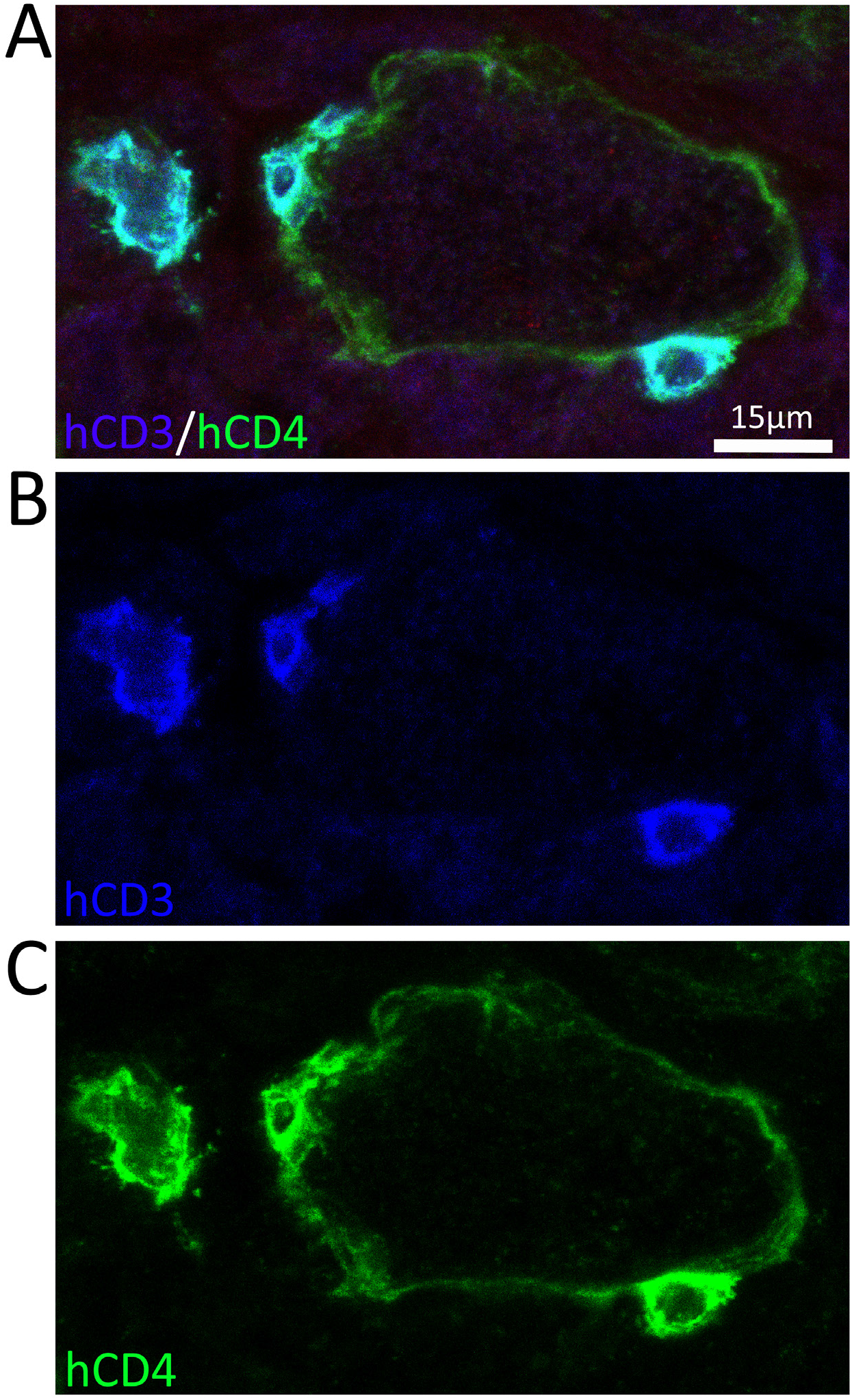
Combined (A) and Isolated (B,C) fluorescent channels from confocal image used in Figure 6E demonstrating human CD3 (B) and human CD4 (C) immunolabeling in the lesion epicenter 35 dpi in a hNSG mouse injured at 4-months post-engraftment. The co-localization of hCD3 and hCD4 on small cells confirms their T cell identity. In this example, two human T cells are in direct contact with a human CD4^+^ macrophage. Note the absence of hCD3 labeling on the human macrophage. A large, isolated T cell with immunoblast-like morphology is also identified.

## Works Cited

Ankeny DP, Guan Z, Popovich PG. 2009. B cells produce pathogenic antibodies and impair recovery after spinal cord injury in mice. J Clin Invest 119:2990–9. doi:10.1172/JCI39780

Audigé A, Rochat M-A, Li D, Ivic S, Fahrny A, Muller CKS, Gers-Huber G, Myburgh R, Bredl S, Schlaepfer E, Scherrer AU, Kuster SP, Speck RF. 2017. Long-term leukocyte reconstitution in NSG mice transplanted with human cord blood hematopoietic stem and progenitor cells. BMC Immunol 18:28. doi:10.1186/s12865-017-0209-9

Baba T, Ishizu A, Iwasaki S, Suzuki A, Tomaru U, Ikeda H, Yoshiki T, Kasahara M. 2006. CD4 /CD8 macrophages infiltrating at inflammatory sites: a population of monocytes/macrophages with a cytotoxic phenotype. Blood 107:2004–2012. doi:10.1182/blood-2005-06-2345

Basso DM, Fisher LC, Anderson AJ, Jakeman LBYNB, Tigue DMMC, Popovich PG, McTigue DM, Popovich PG. 2006. Basso Mouse Scale for locomotion detects differences in recovery after spinal cord injury in five common mouse strains. J Neurotrauma 23:635–659. doi:10.1089/neu.2006.23.635

Beck KD, Nguyen HX, Galvan MD, Salazar DL, Woodruff TM, Anderson AJ. 2010. Quantitative analysis of cellular inflammation after traumatic spinal cord injury: Evidence for a multiphasic inflammatory response in the acute to chronic environment. Brain 133:433–447. doi:10.1093/brain/awp322

Blomster L V., Brennan FH, Lao HW, Harle DW, Harvey AR, Ruitenberg MJ. 2013. Mobilisation of the splenic monocyte reservoir and peripheral CX3CR1 deficiency adversely affects recovery from spinal cord injury. Exp Neurol 247:226–240. doi:10.1016/j.expneurol.2013.05.002

Boddaert J, Bielen K, Manocha E, Yperzeele L, Cras P, Pirici D, Kumar-Singh S. 2018. CD8 signaling in microglia/macrophage M1 polarization in a rat model of cerebral ischemia. doi:10.1371/journal.pone.0186937

Brennan FH, Popovich PG. 2018. Emerging targets for reprograming the immune response to promote repair and recovery of function after spinal cord injury. Curr Opin Neurol. doi:10.1097/WCO.0000000000000550

Carpenter RS, Kigerl KA, Marbourg JM, Gaudet AD, Huey D, Niewiesk S, Popovich PG. 2015. Traumatic spinal cord injury in mice with human immune systems. Exp Neurol 271:432–444. doi:10.1016/j.expneurol.2015.07.011

Cheng L, Zhang Z, Li G, Li F, Wang L, Zhang L, Zurawski SM, Zurawski G, Levy Y, Su L. 2017. Human innate responses and adjuvant activity of TLR ligands in vivo in mice reconstituted with a human immune system. Vaccine 35:6143–6153. doi:10.1016/j.vaccine.2017.09.052

Clayton JA. 2016. Studying both sexes: a guiding principle for biomedicine. FASEB J 30:519–524. doi:10.1096/fj.15-279554

Coughlan AM, Freeley SJ, Robson MG. 2012. Humanised mice have functional human neutrophils. J Immunol Methods 385:96–104. doi:10.1016/j.jim.2012.08.005

Cravens PD, Melkus MW, Padgett-Thomas A, Islas-Ohlmayer M, Del P Martin M, Garcia JV. 2005. Development and activation of human dendritic cells in vivo in a xenograft model of human hematopoiesis. Stem Cells 23:264–278. doi:10.1634/stemcells.2004-0116

Crocker PR, Jefferies WA, Clark SJ, Chung LP, Gordon S. 1987. Species heterogeneity in macrophage expression of the CD4 antigen. J Exp Med 166:613–8. doi:10.1084/JEM.166.2.613

Danner R, Chaudhari SN, Rosenberger J, Surls J, Richie TL, Brumeanu TD, Casares S. 2011. Expression of HLA class II molecules in humanized NOD.Rag1KO.IL2RgcKO mice is critical for development and function of human T and B cells. PLoS One 6:e19826. doi:10.1371/journal.pone.0019826

Dykstra C, Lee AJ, Lusty EJ, Shenouda MM, Shafai M, Vahedi F, Chew M V., Collins S, Ashkar AA. 2016. Reconstitution of immune cell in liver and lymph node of adult- and newborn-engrafted humanized mice. BMC Immunol 17:18. doi:10.1186/s12865-016-0157-9

Esashi E, Sekiguchi T, Ito H, Koyasu S, Miyajima A. 2003. Cutting Edge: A possible role for CD4+ thymic macrophages as professional scavengers of apoptotic thymocytes. J Immunol 171:2773–7. doi:10.4049/JIMMUNOL.171.6.2773

Fleming JC, Norenberg MD, Ramsay DA, Dekaban GA, Marcillo AE, Saenz AD, Pasquale-Styles M, Dietrich WD, Weaver LC. 2006. The cellular inflammatory response in human spinal cords after injury. Brain 129:3249–3269. doi:10.1093/brain/awl296

Fooksman DR, Vardhana S, Vasiliver-Shamis G, Liese J, Blair DA, Waite J, Sacristán C, Victora GD, Zanin-Zhorov A, Dustin ML. 2010. Functional Anatomy of T Cell Activation and Synapse Formation. Annu Rev Immunol 28:79–105. doi:10.1146/annurev-immunol-030409-101308

Gensel JC, Nakamura S, Guan Z, van Rooijen N, Ankeny DP, Popovich PG. 2009. Macrophages Promote Axon Regeneration with Concurrent Neurotoxicity. J Neurosci 29:3956–3968. doi:10.1523/JNEUROSCI.3992-08.2009

Gibbings D, Befus AD. 2009. CD4 and CD8: an inside-out coreceptor model for innate immune cells. J Leukoc Biol 86:251–259. doi:10.1189/jlb.0109040

Gibbings DJ, Marcet-Palacios M, Sekar Y, Ng MC, Befus AD. 2007. CD8α is expressed by human monocytes and enhances FcγR-dependent responses. BMC Immunol 8:12. doi:10.1186/1471-2172-8-12

Gille C, Orlikowsky TW, Spring B, Hartwig UF, Wilhelm A, Wirth A, Goecke B, Handgretinger R, Poets CF, André MC. 2012. Monocytes derived from humanized neonatal NOD/SCID/IL2Rγnullmice are phenotypically immature and exhibit functional impairments. Hum Immunol 73:346–354. doi:10.1016/j.humimm.2012.01.006

Goldstein EZ, Church JS, Hesp ZC, Popovich PG, McTigue DM. 2016. A silver lining of neuroinflammation: Beneficial effects on myelination. Exp Neurol 283:550–559. doi:10.1016/j.expneurol.2016.05.001

Halkias J, Yen B, Taylor KT, Reinhartz O, Winoto A, Robey EAEAE a, Melichar HJ. 2015. Conserved and divergent aspects of human T cell development and migration in humanized mice. Immunol Cell Biol 93:1–11. doi:10.1038/icb.2015.38

Hauben Ehud, Butovsky O, Nevo U, Yoles E, Moalem G, Agranov E, Mor F, Leibowitz-Amit R, Pevsner E, Akselrod S, Neeman M, Cohen IR, Schwartz M. 2000. Passive or Active Immunization with Myelin Basic Protein Promotes Recovery from Spinal Cord Contusion. J Neurosci 20:6421–6430.

Hauben E., Nevo U, Yoles E, Moalem G, Agranov E, Mor F, Akselrod S, Neeman M, Cohen IR, Schwartz M. 2000. Autoimmune T cells as potential neuroprotective therapy for spinal cord injury. Lancet 355:286–287. doi: 10.1016/S0140-6736(99)05140-5

Hiraki K, Park IK, Kohyama K, Matsumoto Y. 2009. Characterization of CD8-positive macrophages infiltrating the central nervous system of rats with chronic autoimmune encephalomyelitis. J Neurosci Res 87:1175–1184. doi:10.1002/jnr.21924

Holladay SD, Smialowicz RJ. 2000. Development of the murine and human immune system: differential effects of immunotoxicants depend on time of exposure. Environ Health Perspect 108:463–473. doi:10.1289/ehp.00108s3463

Huey DD, Niewiesk S. 2018. Production of Humanized Mice Through Stem Cell Transfer. Curr Protoc Mouse Biol 8:17–27. doi:10.1002/cpmo.38

Ikeda H, Iwasaki S, Suzuki A, Maruoka T, Ishizu A, Baba T, Yoshiki T, Kasahara M, Tomaru U. 2014. Rat CD4+CD8+ Macrophages Kill Tumor Cells through an NKG2D- and Granzyme/Perforin-Dependent Mechanism. J Immunol 180:2999–3006. doi:10.4049/jimmunol.180.5.2999

Ishikawa F, Yasukawa M, Lyons B, Yoshida S, Miyamoto T, Yoshimoto G, Watanabe T, Akashi K, Shultz LD, Harada M. 2005. Development of functional human blood and immune systems in NOD/SCID/IL2 receptor γ chainnull mice. Blood 106:1565–1573. doi:10.1182/blood-2005-02-0516

Jones TB. 2014. Lymphocytes and autoimmunity after spinal cord injury. Exp Neurol 258:78–90. doi:10.1016/j.expneurol.2014.03.003

Jones TB, Ankeny DP, Guan Z, McGaughy V, Fisher LC, Basso DM, Popovich PG. 2004. Passive or Active Immunization with Myelin Basic Protein Impairs Neurological Function and Exacerbates Neuropathology after Spinal Cord Injury in Rats. J Neurosci 24:3752–3761. doi:10.1523/JNEUROSCI.0406-04.2004

Jones TB, Basso DM, Sodhi A, Pan JZ, Hart RP, MacCallum RC, Lee S, Whitacre CC, Popovich PG. 2002. Pathological CNS autoimmune disease triggered by traumatic spinal cord injury: implications for autoimmune vaccine therapy. J Neurosci 22:2690–2700. doi:20026267

Jones TB, Hart RP, Popovich PG. 2005. Molecular Control of Physiological and Pathological T-Cell Recruitment after Mouse Spinal Cord Injury. J Neurosci 25:6576–6583. doi:10.1523/JNEUROSCI.0305-05.2005

Kigerl KA, Gensel JC, Ankeny DP, Alexander JK, Donnelly DJ, Popovich PG. 2009. Identification of two distinct macrophage subsets with divergent effects causing either neurotoxicity or regeneration in the injured mouse spinal cord. J Neurosci 29:13435–44. doi:10.1523/JNEUROSCI.3257-09.2009

Kigerl KA, McGaughy VM, Popovich PG. 2006. Comparative analysis of lesion development and intraspinal inflammation in four strains of mice following spinal contusion injury. J Comp Neurol 494:578–594. doi:10.1002/cne.20827

Knibbe-Hollinger JS, Fields NR, Chaudoin TR, Epstein AA, Makarov E, Akhter SP, Gorantla S, Bonasera SJ, Gendelman HE, Poluektova LY. 2015. Influence of age, irradiation and humanization on NSG mouse phenotypes. Biol Open 1–10. doi:10.1242/bio.013201

Lang J, Kelly M, Freed BM, McCarter MD, Kedl RM, Torres RM, Pelanda R. 2013. Studies of Lymphocyte Reconstitution in a Humanized Mouse Model Reveal a Requirement of T Cells for Human B Cell Maturation. J Immunol 190:2090–2101. doi:10.4049/jimmunol.1202810

Liu Z, Zhang H, Xia H, Wang Baocheng, Zhang R, Zeng Q, Guo L, Shen K, Wang BaTa, Zhong Y, Li Z, Sun G. 2019. CD8 T cell-derived perforin aggravates secondary spinal cord injury through destroying the blood-spinal cord barrier. Biochem Biophys Res Commun. doi:10.1016/J.BBRC.2019.03.002

Mestas J, Hughes CCW. 2004. Of mice and not men: differences between mouse and human immunology. J Immunol 172:2731–2738. doi:10.4049/jimmunol.172.5.2731

Milićević NM, Nohroudi K, Milićević Z, Hedrich H-J, Westermann J. 2005. T cells are required for the peripheral phase of B-cell maturation. Immunology 116:308–17. doi:10.1111/j.1365-2567.2005.02226.x

Miller PH, Cheung AMS, Beer PA, Knapp DJHF, Dhillon K, Rabu G, Rostamirad S, Humphries RK, Eaves CJ. 2013. Enhanced normal short-term human myelopoiesis in mice engineered to express human-specific myeloid growth factors. Blood 121:e1–4. doi:10.1182/blood-2012-09-456566

Misharin A V, Haines GK, Rose S, Gierut AK, Hotchkiss RS, Perlman H. 2012. Development of a new humanized mouse model to study acute inflammatory arthritis. J Transl Med 10:190. doi:10.1186/1479-5876-10-190

Noble BT, Brennan FH, Popovich PG. 2018. The spleen as a neuroimmune interface after spinal cord injury. J Neuroimmunol. doi:10.1016/j.jneuroim.2018.05.007

O’Boyle G, Fox CRJ, Walden HR, Willet JDP, Mavin ER, Hine DW, Palmer JM, Barker CE, Lamb C a., Ali S, Kirby J a. 2012. Chemokine receptor CXCR3 agonist prevents human T-cell migration in a humanized model of arthritic inflammation. Proc Natl Acad Sci 109:4598–4603. doi:10.1073/pnas.1118104109

Pearson T, Greiner DL, Shultz LD. 2008. Creation of “humanized” Mice to study human immunity. Curr Protoc Immunol Chapter 15:Unit 15.21. doi:10.1002/0471142735.im1521s81

Popovich PG, Guan Z, McGaughy V, Fisher L, Hickey WF, Basso DM. 2002. The neuropathological and behavioral consequences of intraspinal microglial/macrophage activation. J Neuropathol Exp Neurol 61:623–633.

Popovich PG, Van Rooijen N, Hickey WF, Preidis G, McGaughy V. 2003. Hematogenous macrophages express CD8 and distribute to regions of lesion cavitation after spinal cord injury. Exp Neurol 182:275–287. doi:10.1016/S0014-4886(03)00120-1

Raposo C, Graubardt N, Cohen M, Eitan C, London A, Berkutzki T, Schwartz M. 2014. CNS repair requires both effector and regulatory T cells with distinct temporal and spatial profiles. J Neurosci 34:10141–55. doi:10.1523/JNEUROSCI.0076-14.2014

Rathinam C, Poueymirou WT, Rojas J, Murphy AJ, Valenzuela DM, Yancopoulos GD, Rongvaux A, Eynon EE, Manz MG, Flavell RA. 2011. Efficient differentiation and function of human macrophages in humanized CSF-1 mice. Blood 118:3119–3128. doi:10.1182/blood-2010-12-326926

Rodewohl A, Scholbach J, Leichsenring A, Köberle M, Lange F. 2017. Age-dependent cellular reactions of the human immune system of humanized NOD scid gamma mice on LPS stimulus. Innate Immun 23:258–275. doi:10.1177/1753425917690814

Rongvaux A, Willinger T, Martinek J, Strowig T, Gearty S V, Teichmann LL, Saito Y, Marches F, Halene S, Palucka a K, Manz MG, Flavell R a. 2014. Development and function of human innate immune cells in a humanized mouse model. Nat Biotechnol 32:364–72. doi:10.1038/nbt.2858

Russell A, Malik S, Litzow M, Gastineau D, Roy V, Zubair AC. 2015. Dual roles of autologous CD8+ T cells in hematopoietic progenitor cell mobilization and engraftment. Transfusion 55:1758–1765. doi:10.1111/trf.13073

Saito Y, Ellegast JM, Rafiei A, Song Y, Kull D, Heikenwalder M, Rongvaux A, Halene S, Flavell RA, Manz MG. 2016. Peripheral blood CD34+ cells efficiently engraft human cytokine knock-in mice. Blood 128:1829–1833. doi:10.1182/blood-2015-10-676452

Satzer D, Miller C, Maxon J, Voth J, DiBartolomeo C, Mahoney R, Dutton JR, Low WC, Parr AM. 2015. T cell deficiency in spinal cord injury: altered locomotor recovery and whole-genome transcriptional analysis. BMC Neurosci 16:74. doi:10.1186/s12868-015-0212-0

Schürch CM, Riether C, Ochsenbein AF. 2014. Cytotoxic CD8+ T cells stimulate hematopoietic progenitors by promoting cytokine release from bone marrow mesenchymal stromal cells. Cell Stem Cell 14:460–472. doi:10.1016/j.stem.2014.01.002

Schwartz M. 2001. Protective autoimmunity as a T-cell response to central nervous system trauma: Prospects for therapeutic vaccines. Prog Neurobiol 65:489–496. doi:10.1016/S0301-0082(01)00009-0

Schwartz M, Kipnis J. 2001. Protective autoimmunity: Regulation and prospects for vaccination after brain and spinal cord injuries. Trends Mol Med 7:252–258. doi:10.1016/S1471-4914(01)01993-1

Schwartz M, Moalem G. 2001. Beneficial immune activity after CNS injury: Prospects for vaccination. J Neuroimmunol 113:185–192. doi:10.1016/S0165-5728(00)00447-1

Shechter R, Schwartz M. 2013. Harnessing monocyte-derived macrophages to control central nervous system pathologies: No longer “if” but “how.” J Pathol 229:332–346. doi:10.1002/path.4106

Shultz LD, Brehm M a, Garcia-Martinez JV, Greiner DL. 2012. Humanized mice for immune system investigation: progress, promise and challenges. Nat Rev Immunol 12:786–98. doi:10.1038/nri3311

Shultz LD, Brehm MA, Bavari S, Greiner DL. 2011. Humanized mice as a preclinical tool for infectious disease and biomedical research. Ann N Y Acad Sci 1245:50–54. doi:10.1111/j.1749-6632.2011.06310.x

Shultz LD, Ishikawa F, Greiner DL. 2007. Humanized mice in translational biomedical research. Nat Rev Immunol 7:118–30. doi:10.1038/nri2017

Shultz LD, Saito Y, Najima Y, Tanaka S, Ochi T, Tomizawa M, Doi T, Sone A, Suzuki N, Fujiwara H, Yasukawa M, Ishikawa F. 2010. Generation of functional human T-cell subsets with HLA-restricted immune responses in HLA class I expressing NOD/SCID/IL2r null humanized mice. Proc Natl Acad Sci 107:13022–13027. doi:10.1073/pnas.1000475107

Sosa JM, Huber DE, Welk B, Fraser HL. 2014. Development and application of MIPAR™: a novel software package for two- and three-dimensional microstructural characterization. Integr Mater Manuf Innov 3:10. doi:10.1186/2193-9772-3-10

Sroga JM, Jones TB, Kigerl K a, McGaughy VM, Popovich PG. 2003. Rats and mice exhibit distinct inflammatory reactions after spinal cord injury. J Comp Neurol 462:223–40. doi:10.1002/cne.10736

Strowig T, Chijioke O, Carrega P, Arrey F, Meixlsperger S, Rämer PC, Ferlazzo G, Münz C. 2010. Human NK cells of mice with reconstituted human immune system components require preactivation to acquire functional competence. Blood 116:4158–4167. doi:10.1182/blood-2010-02-270678

Sun G, Yang S, Cao G, Wang Q, Hao J, Wen Q, Li Z, So K-F, Liu Z, Zhou S, Zhao Y, Yang H, Zhou L, Yin Z. 2017. γδ T cells provide the early source of IFN-γ to aggravate lesions in spinal cord injury. J Exp Med 215:521–535. doi:10.1084/jem.20170686

Szabo G, Miller CL, Kodys K. 1990. Antigen presentation by the CD4 positive monocyte subset. J Leukoc Biol 47:111–120. doi:10.1002/jlb.47.2.111

Takagi S, Saito Y, Hijikata A, Tanaka S, Watanabe T, Hasegawa T, Mochizuki S, Kunisawa J, Kiyono H, Koseki H, Ohara O, Saito T, Taniguchi S, Shultz LD, Ishikawa F. 2012. Membrane-bound human SCF/KL promotes in vivo human hematopoietic engraftment and myeloid differentiation. Blood 119:2768–2777. doi:10.1182/blood-2011-05-353201

Tanaka S, Saito Y, Kunisawa J, Kurashima Y, Wake T, Suzuki N, Shultz LD, Kiyono H, Ishikawa F. 2012. Development of Mature and Functional Human Myeloid Subsets in Hematopoietic Stem Cell-Engrafted NOD/SCID/IL2r KO Mice. J Immunol 188:6145–6155. doi:10.4049/jimmunol.1103660

Walsh JT, Hendrix S, Boato F, Smirnov I, Zheng J, Lukens JR, Gadani S, Hechler D, Gölz G, Rosenberger K, Kammertöns T, Vogt J, Vogelaar C, Siffrin V, Radjavi A, Fernandez-Castaneda A, Gaultier A, Gold R, Kanneganti T-D, Nitsch R, Zipp F, Kipnis J. 2015. MHCII-independent CD4+ T cells protect injured CNS neurons via IL-4. J Clin Invest 125:699–714. doi:10.1172/JCI76210

Walsh JT, Zheng J, Smirnov I, Lorenz U, Tung K, Kipnis J. 2014. Regulatory T cells in central nervous system injury: a double-edged sword. J Immunol 193:5013–5022. doi:10.4049/jimmunol.1302401

Wetmore A, Herlihy M, Hosur V, Leif J, Greiner DL, Racki WJ, Gott B, Shultz LD, Brehm MA, Burzenski L, Ignotz R, Dunn R. 2012. Engraftment of human HSCs in nonirradiated newborn NOD-scid IL2r null mice is enhanced by transgenic expression of membrane-bound human SCF. Blood 119:2778–2788. doi:10.1182/blood-2011-05-353243

Wu Y, Lin YH, Shi LL, Yao ZF, Xie XM, Jiang ZS, Tang J, Hu JG, Lü HZ. 2017. Temporal kinetics of CD8+CD28+and CD8+CD28-T lymphocytes in the injured rat spinal cord. J Neurosci Res 95:1666–1676. doi:10.1002/jnr.23993

Wunderlich M, Chou FS, Sexton C, Presicce P, Chougnet CA, Aliberti J, Mulloy JC. 2018. Improved multilineage human hematopoietic reconstitution and function in NSGS mice. PLoS One 13:e0209034. doi:10.1371/journal.pone.0209034

Yoshihara S, Li Y, Xia J, Danzl N, Sykes M, Yang Y-G. 2019. Posttransplant Hemophagocytic Lymphohistiocytosis Driven by Myeloid Cytokines and Vicious Cycles of T-Cell and Macrophage Activation in Humanized Mice. Front Immunol 10:186. doi:10.3389/fimmu.2019.00186

Zangerle-Murray T, Tamoutounour S, Houston SA, Barbera TA, Allen JE, Bridgeman HM, Konkel JE, Shaw TN, Wang P, Strangward P, Wemyss K, Ridley AJL, Grainger JR. 2018. Tissue-resident macrophages in the intestine are long lived and defined by Tim-4 and CD4 expression. J Exp Med 215:1507–1518. doi:10.1084/jem.20180019

Zayoud M, El Malki K, Frauenknecht K, Trinschek B, Kloos L, Karram K, Wanke F, Georgescu J, Hartwig UF, Sommer C, Jonuleit H, Waisman A, Kurschus FC. 2013. Subclinical CNS inflammation as response to a myelin antigen in humanized mice. J Neuroimmune Pharmacol 8:1037–1047. doi:10.1007/s11481-013-9466-4

Zhen A, Zack JA, Kasparian S, Levin BR, Kitchen SG, Krutzik SR. 2014. CD4 Ligation on Human Blood Monocytes Triggers Macrophage Differentiation and Enhances HIV Infection. J Virol 88:9934–9946. doi:10.1128/jvi.00616-14

